# Beat processing in newborn infants cannot be explained by statistical learning based on transition probabilities

**DOI:** 10.1101/2022.12.20.521245

**Authors:** Gábor P. Háden, Fleur L. Bouwer, Henkjan Honing, István Winkler

## Abstract

Newborn infants have been shown to extract temporal regularities from sound sequences, both in the form of learning regular sequential properties, and extracting periodicity in the input, commonly referred to as a beat. However, these two types of regularities are often indistinguishable in isochronous sequences, as both statistical learning and beat perception can be elicited by the regular alternation of accented and unaccented sounds. Here, we manipulated the isochrony of sound sequences in order to disentangle statistical learning from beat perception in sleeping newborn infants in an EEG experiment, as previously done in adults and macaque monkeys. We used a binary accented sequence that induces a beat when presented with isochronous timing, but not when presented with randomly jittered timing. We compared mismatch responses to infrequent deviants falling on either accented or unaccented (i.e., odd and even) positions. Results showed a clear difference between metrical positions in the isochronous sequence, but not in the equivalent jittered sequence. This suggests that beat processing is present in newborns. However, the current paradigm did not show effects of statistical learning, despite previous evidence for this ability in newborns. These results show that statistical learning does not explain beat processing in newborn infants.

**Research highlights:** Sleeping newborns process musical beat.

Transition probabilities are not enough to explain beat perception in newborn infants.

No evidence of statistical learning (based on transition probabilities) without isochronous stimulation in newborns.

Results converge with previous evidence on beat perception of newborn infants.

## Introduction

Processing sound sequences is crucial for both music and speech perception (Patel, 2008; Kotz et al., 2018). Studying the development of sound processing in infants can provide insights into the bootstrapping of speech and music acquisition during a period in which learning and brain maturation go hand in hand (Finlay & Uchiyama, 2020). In sound sequences, several types of structure can be discerned that contribute to efficient processing, including regularities in the temporal aspects of a sequence (i.e., the ‘beat’), regularities in order of events (i.e., statistical regularities), and hierarchical temporal stress patterns (i.e., the meter; Langus, Mehler, & Nespor, 2017). While both the perception of temporal (Winkler et al., 2009) and statistical (Bosseler et al., 2016) regularities have been suggested to be present in neonates, the relationship between these remains unclear, since sequences used to probe statistical regularities often also contain temporal regularities and vice versa (Bouwer et al., 2014). Here we aim to disentangle the ability of neonates to detect regularities in time (the beat or pulse), and regularities in order (statistical learning).

Temporal structure in rhythm often comes in the form of a regular beat: a series of perceived regularly recurring salient events (Honing & Bouwer, 2019; Honing, 2012; Cooper & Meyer, 1960). The ability to synchronize and coordinate movement with a regular beat is called rhythmic entrainment. This coordination is thought to result from the coupling of internal low frequency oscillations in auditory and motor areas with the external rhythmic signal (neural entrainment; see Large, Herrera & Valesco, 2015). Beat-based timing has been suggested to be somewhat separate from other timing processes, like the perception of (a sequence of) single absolute temporal intervals (Honing & Merchant, 2014; Teki et al., 2011; Breska & Deouell, 2017; Bouwer, Honing & Slagter, 2020). In humans, we have previously suggested that beat perception is already functional at birth (Winkler et al., 2009). However, the results of this study could have been biased by comparing between responses to acoustically different sounds and contexts (detailed in Bouwer et al., 2014). Hence, the presence of beat perception at birth remains to be confirmed. (Note that we will refer to the detection of beat as shown by brain responses as “beat perception” to keep the terminology compatible with previous literature, although infants cannot describe their experience, making this term incorrect in the literal sense.)

Specifically, beat perception needs to be dissociated from the detection of statistical regularities in the order of events, such as transitional/conditional probabilities of item succession, as was described for language-like (Saffran, Newport & Aslin, 1996) and in tonal stimuli (Saffran, et al., 1999). Statistical learning in its narrowest sense is the extraction of these regularities from the order of the elements of the input without explicit feedback or even awareness after prolonged exposure (Conway, 2020). Statistical learning is thought to be a domain general mechanism manifest in different domains, including audition (Saffran, Newport & Aslin, 1996), music (Saffran, et al., 1999, Pearce, 2018), and vision (Fiser & Aslin, 2001; Duncan & Theeuwes, 2020), and most prominently in language. There is evidence that statistical learning is already functional at birth (Bulf, Johnson & Valenza, 2011; Bosseler et al., 2016, Teinonen, et al., 2009). However, there is an ongoing debate whether statistical learning in children is an unchanging domain general mechanism or that it shows developmental changes that differ between the visual and the auditory modality (e.g., Saffran et al., 1997; Raviv & Arnon, 2018; reviewed in Conway, 2020).

Understanding the processing of sound sequences in infants requires one to dissociate the processing of the beat, absolute temporal intervals, and statistical regularities in the order of sounds. Firstly, to show beat perception, the rhythmic stimulus needs to be acoustically and/or temporally varied, so that these can be distinguished from interval-based perception (Bouwer et al., 2016, 2020; Honing et al., 2018). Secondly, while the presence of acoustic variation in a sequence (e.g., differences in pitch or intensity between events) can aid the detection of a beat, especially in musical novices (Bouwer, Burgoyne, Odijk, Honing & Grahn, 2018), it also creates possible confounds when probing beat perception, as differences in responses related to the temporal structure (e.g., different responses to events on and off the beat) may in fact be caused by differences in acoustic and sequential order properties when using acoustically rich and varied rhythmic sequences (Bouwer et al., 2014). Thus, to show beat perception, it must be carefully dissociated from learning transitional probabilities.

To disentangle beat perception from statistical learning and the perception of interval-based temporal structure (e.g., learning an absolute interval), an auditory oddball paradigm was proposed by Bouwer et al. (2016; also used in Honing et al., 2018). The paradigm employs a rhythmic sequence made up of a pattern of loud (on all odd, “beat” positions and, a small portion of even, “offbeat” positions) and soft (in most even, “offbeat” positions) percussive sounds (Figure 1A), such that the acoustic stimulus could induce a simple binary metrical structure (“duple meter”). The presence of timbre and intensity differences arguably creates an ecological way of inducing a beat (Ladinig et al., 2009). Within the context of this clearly beat-inducing sequence of alternating loud and soft sounds, in a proportion of the patterns, the offbeat positions are also filled with loud sounds. These patterns are used to probe beats and offbeats with identical acoustic properties. These rhythmic sequences are presented in two conditions: an isochronous condition, and a jittered condition (Figure 1B). In the isochronous condition, sounds are presented with a constant inter-onset interval (IOI), allowing a beat to be induced (i.e., one metrical level of a duple meter). In the jittered condition, the IOIs are irregular, thus disabling the perception of a regular beat. However, the jittered sequences still contain the same order-based statistical regularity (alternation) between the louder and softer sounds as the isochronous ones.

**Figure 1.**
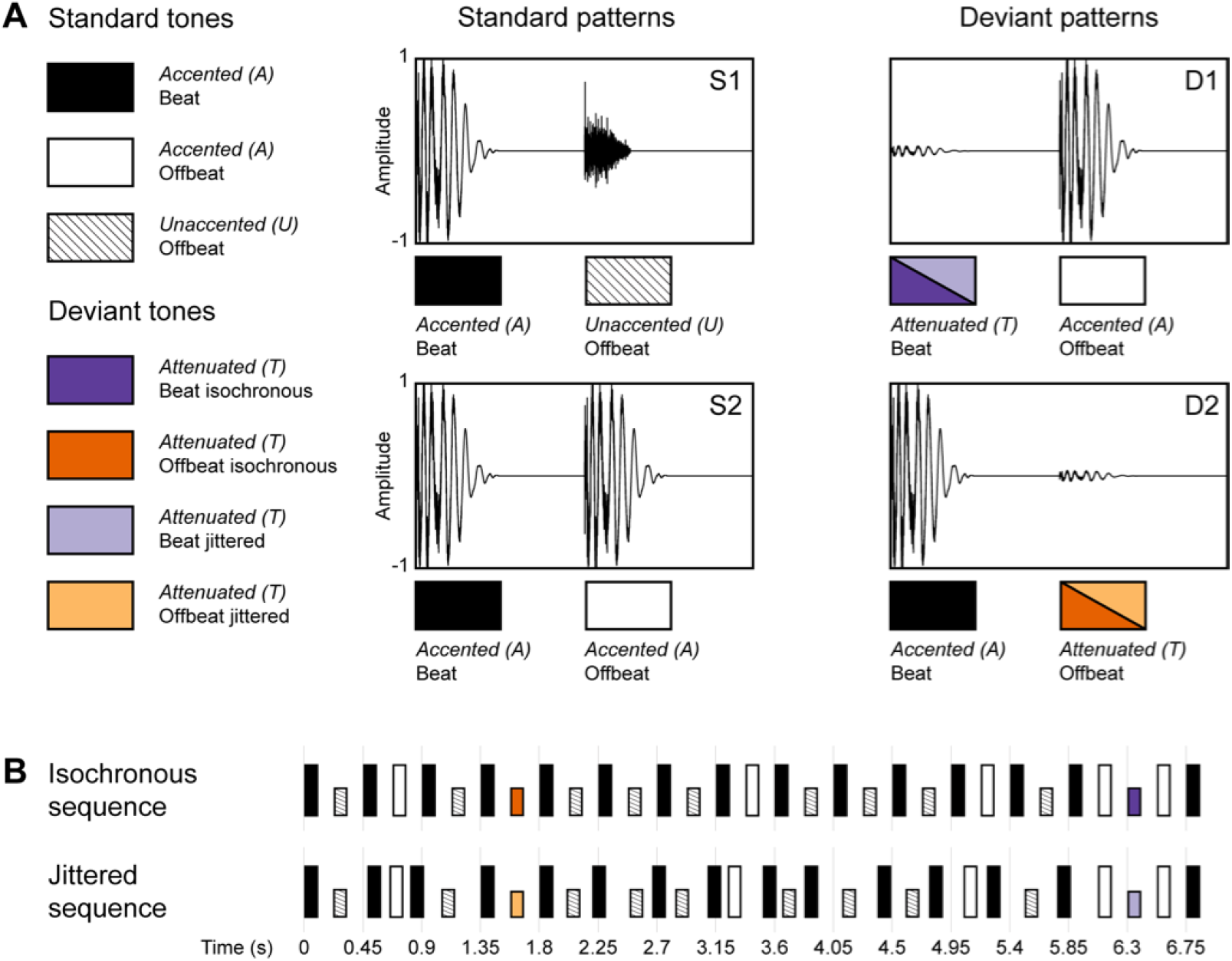
(color) Schematic diagram of the rhythmic stimulus patterns used in the experiment. (A) The two standard (S1 and S2) and two deviant patterns (D1 and D2) are made up of three different sounds (A=accented, U=unaccented, and T=attenuated). An accented sound could occur either on the beat or offbeat, an unaccented sound was restricted to the offbeat position. Attenuated sounds were used as deviants in both positions (beat and offbeat) and conditions (isochronous and jittered). (B) Standard and deviant sound patterns were concatenated into a single rhythmic stream in a random order (see main text for details). Sequences in the isochronous condition had an inter-onset interval (IOI) of 225 ms, in the jittered condition these were randomly chosen from the range 150 to 300 ms using a uniform distribution. Deviants were always preceded and followed by an accented sound, with a fixed IOI of 225 ms in both conditions. Note that while we use the labels Beat and Offbeat in both conditions, these are only perceptually valid in the isochronous condition. (Adapted from Honing et al., 2018).

As such, the jittered condition serves as a control for statistical learning of the succession of sounds with different timbres.

Bouwer et al. (2016) presented these sequences to adults, and measured event-related potentials (ERPs) to rare, unexpected intensity decrements (i.e., deviants) to assess the formation of beat-based expectations. If the prediction of incoming stimuli based on previous stimuli fails, that is, the regularity extracted from the sound sequence is violated, the mismatch negativity (MMN) ERP component is elicited in adults (for a recent review, see Fitzgerald & Todd, 2020). The amplitude of the MMN increases together with the specificity of the prediction (Southwell & Chait, 2018) and the amount of deviation from the predicted sound (e.g., Novitski et al., 2004). Bouwer et al. (2016) found larger MMN responses for deviants in odd (beat) than even (offbeat) positions, even in the jittered condition, indicative of statistical learning that resulted in different predictions for beat and offbeat positions. The MMN responses were much larger in the isochronous than in the jittered condition, which the authors took as support for the presence of beat perception. Arguably, beat perception contributed to differences between positions only in the isochronous, but not the jittered condition. Moreover, in both positions, the P3a (an ERP component indexing post-processing following the detection of an unattended regularity violation; for a review, see Polich, 2007) was larger in the isochronous than the jittered condition, indicating that the isochrony of the sequence increased the amount of processing needed after a regularity violation.

A mismatch response (MMR) similar to MMN can be measured in young infants in response to regularity violations, including sleeping neonates (Alho et al., 1990, Háden et al., 2016). In our previous study in newborn infants (Winkler et al., 2009) stimulus omissions (Háden et al., 2015) at different metrical positions of a repetitive rhythmic sequence were used to test beat and meter processing. However, as was noted before, the different metrical positions had different acoustic and sequential properties. This makes it possible that differences in omission MMR responses found for different metrical positions were based on learning the statistical properties of the order of different tones, that is, by statistical learning (Bouwer et al., 2014).

Here, we presented a variant of Bouwer et al.’s (2016) stimulus paradigm to sleeping newborn infants to separate the effects of beat perception from those of statistical learning (Figure 1) in order to provide converging evidence to the notion that newborns can perceive a beat in rhythm, as suggested in our previous study (Winkler et al, 2009), while controlling for the effects of statistical learning. Response differences (MMR) between rare deviant and corresponding standard sounds were calculated at beat (odd) and offbeat (even) positions, separately for the isochronous and the jittered sequences (note that we refer to odd and even positions as “beat” and “offbeat” even for the jittered sequences, to make the terminology more consistent). In line with the effects in adults, firstly we expected newborns to learn the statistically predictable alternation between louder and softer sounds. Since beat perception does not occur in the jittered condition and the MMR is calculated by comparing the responses to sounds with identical acoustic properties and context (i.e., no acoustic difference between the tones in the compared nor in the preceding position), differences in the MMR responses between beat and offbeat positions should result from learning the sequential statistical regularities of the sound sequence (i.e., the alternation). Secondly, we expected to find evidence for beat perception, to corroborate our previous results (c.f. Bouwer et al., 2016 as well as Winkler et al., 2009). Based on Bouwer et al.’s (2016) results, beat perception should make the MMR difference between beat and offbeat position larger in the isochronous than in the jittered condition, as it would additionally contribute to the beat-offbeat difference.

## Material and methods

### Participants

EEG was recorded from 31 (12 male) healthy, full-term newborn infants during day 0–6 (mean=2.06, SD=1.01) postpartum. The mean gestational age was 39.31 weeks (SD=0.87), mean birth weight was 3319 g (SD=311.80). All participants had an Apgar score of 9/10 (1 min./5 min.). Due to excessive electrical artifacts, 4 participants with less than 50% of the trials retained for any of the tested stimuli were excluded from the analysis. The final sample consisted of 27 (12 male) participants. Infants were predominantly asleep during the recordings, in quiet sleep 87% of the time and in active sleep 11% of the time (Anders, Emde & Parmelee, 1971). Informed consent was obtained from the mother or both parents. The experiment was carried out in a dedicated experimental room.

### Stimuli

The stimuli were identical to those used in Bouwer et al. (2016) and Honing et al. (2018), and the following description was adapted from Honing et al. (2018). See Figure 1 for a schematic depiction of the rhythmic stimulus patterns used, Figure 2 for a depiction of the stimuli generation as a transition network, and Supplementary Materials for audio examples of the two conditions.

**Figure 2.**
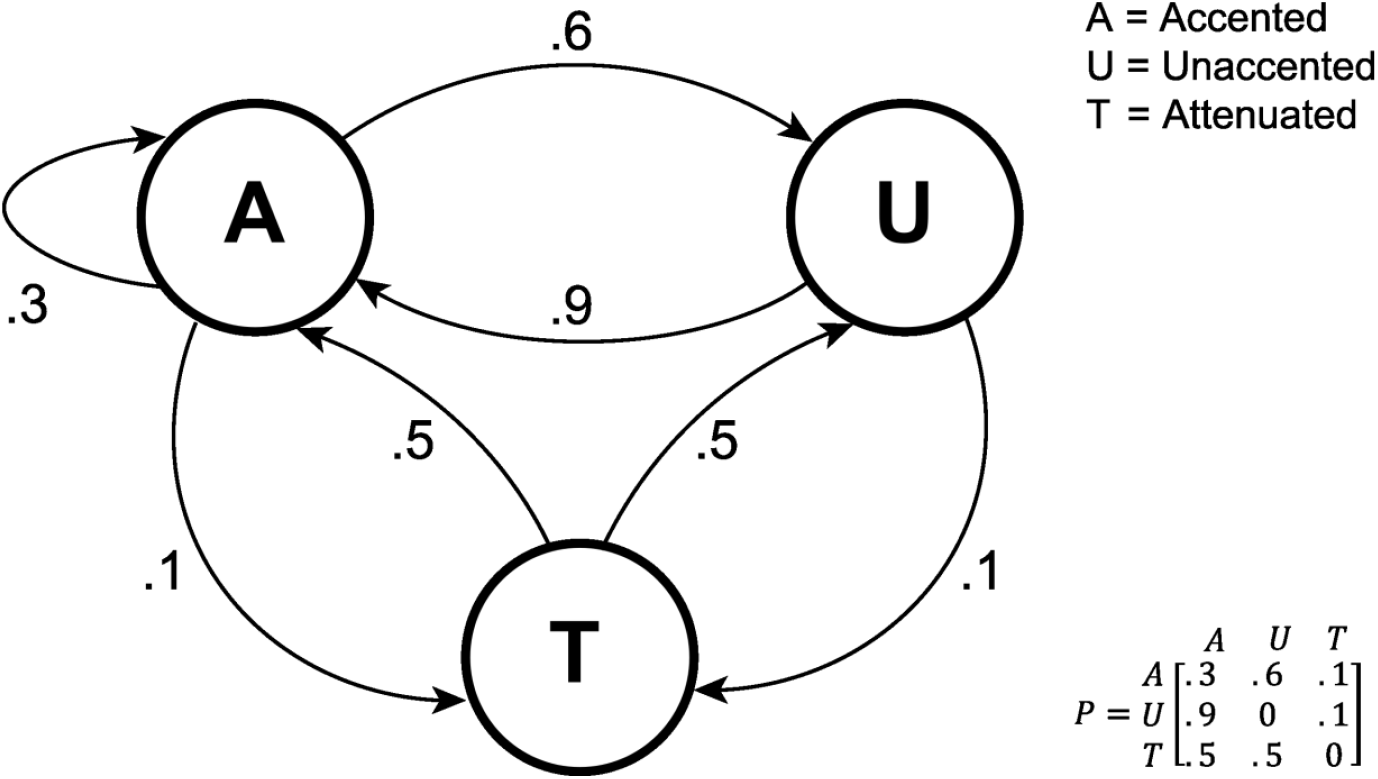
Rhythmic sequence generation visualized as a transition network, showing the transition probabilities between the individual sound elements A, U and T. Note that this representation is a simplification as it does not include certain constraints on the way concatenated patterns can follow each other (see main text). (Adapted from Honing et al., 2018).

To induce a simple binary metrical structure (duple meter) two sounds differing in timbre, intensity and duration were used. The sounds were made with QuickTime’s (Apple, Inc., Cupertino, CA, U.S.A.) drum timbres. The first sound (hereafter Accented or A) consisted of simultaneous sounding of bass drum and hi-hat, whereas the second sound (hereafter Unaccented or U) consisted of a hi-hat sound. Sound lengths were 110 ms and 70 ms, respectively. A third, deviant sound (hereafter Attenuated or T), was created by attenuating the accented sound by 25 dB using Praat software (www.praat.org).

A rhythmic sound sequence was created by concatenating patterns consisting of two sounds. Sixty percent of the patterns consisted of an accented sound followed by an unaccented sound (S1, used to create a context with a clear duple meter) and 30% of the patterns consisted of an accented sound followed by another accented sound (S2, used to create offbeats with an accented sound, to allow deviants on the beat to have the same acoustic context as deviants off the beat). In the remaining 10% of the sequence either an attenuated sound followed an accented sound (5%; D1, deviant in even position, offbeat) or an accented sound followed an attenuated sound (5%; D2, deviant in odd position, beat). Concatenating the four patterns in a random order resulted in accented and unaccented sounds alternating most of the time.

The resulting sequence was used in both the isochronous and jittered conditions. In the isochronous condition, all sounds were presented with a constant inter-onset interval of 225 ms. In this condition, the alternation of accented and unaccented sounds was expected to induce a beat (or duple meter) with an inter-beat interval of 450 ms, which is within the optimal range for beat perception in humans (London, 2002). Sounds in odd positions of the sequence (including deviant D1) can be considered on the beat, while sounds in even positions (including deviant D2) offbeat. In the jittered condition, the inter-onset intervals were randomly distributed between 150 and 300 ms with an average of 225 ms (uniform distribution). This resulted in a sequence with the same order of stimuli as the isochronous one, while not inducing a regular beat (London, 2002; Honing, 2012) (note that D1 and D2 will be referred to as on the beat or offbeat for both conditions, although perceptually this is only true for the isochronous condition).

Sequences were created with the following five constraints. The inter-onset interval just before and after a deviant tone was kept constant at 225ms. A deviant on the beat (D1) was always preceded by an accented sound offbeat (S2). These two constraints made the acoustic and preceding temporal context of the deviants identical between positions and conditions. To optimize the possibility of inducing beat in the isochronous condition, S2 (containing two consecutive accented sounds) was never repeated, and only a maximum of four consecutive S1 patterns was allowed. Finally, at least five standard patterns (S1 and S2) were delivered between two consecutive deviant patterns.

An alternative description of the rhythmic sound sequence generation (at the level of the individual sounds A, U and T) is given in Figure 2. This transition network shows the transition probabilities between the three individual sounds used in both the isochronous and jittered conditions.

### Procedure

Stimuli were presented in 6 blocks with 3 blocks consisting of isochronous and 3 blocks of jittered sequences (1,300 patterns, i.e., 2,600 sound events, per block). One block lasted 9 min and 45 s, and blocks were separated by a silent interval of at least 15 s. Isochronous and jittered blocks were presented in semi-random order, with a maximum of two blocks from the same condition following each other. This resulted in a total session length of about 60 min.

Sounds were presented binaurally using the E-Prime stimulus presentation software (version 1.1.4.1, Psychology Software Tools, Inc., Pittsburgh, PA, USA) via ER-1 headphones (Etymotic Research Inc., Elk Grove Village, IL, USA) connected via sound tubes to self-adhesive ear cups (Sanibel Supply, Middelfart, Denmark) placed over the infants’ ears. The sound intensity was set to a comfortable level for an adult listener. The output of the sound system was hard limited to 80 dB SPL using a purpose built and calibrated circuit between the headphones and computer.

Overall, the design of the study was identical to the unattended condition presented in Bouwer et al. (2016), except that the adult participants watched a silent video with subtitles, whereas infants were mostly asleep (see above.)

### EEG recording and analysis

EEG was recorded with Ag/AgCl electrodes attached to the Fp1, Fp2, Fz, F3, F4, F7, F8, T3, T4, Cz, C3, C4, Pz, P3, P4 locations according to the International 10-20 system, and near to the outer canthus of the right eye. The reference electrode was placed on the tip of the nose and the ground electrode on the forehead. Data was recorded using a direct-coupled amplifier (V-Amp, Brain Products, Munich, Germany) at 24-bit resolution with a sampling rate of 1000 Hz. Signals were off-line filtered between 1.5–30 Hz and epochs of -100–500 ms with respect to sound onset were extracted for each of the D1, D2, S1 (only after S2 to control for acoustic context; for brevity referred to simply as S1 from this point on), and S2 stimuli. The 100-ms pre-stimulus interval served as the baseline for amplitude measurements and illustrations. Due to low signal-to-noise ratio on other electrodes only data from electrodes F3, Fz, F4, C3, Cz, C4, P3, Pz, and P4 were further analyzed. Matlab (version 8.3, Mathworks,Inc., Nantick, MA, USA), EEGLAB (version 2019, Delorme & Makeig, 2004), and ERPLAB (version 7.0, Lopez-Calderon & Luck, 2014) were used for data processing.

Epochs with a voltage change exceeding 100 μV on any channels measured in a moving window (window length = 100 ms, window step = 50 ms) for the entire epoch duration were rejected from the analysis. The mean percentage of accepted trials was 83 (SD=12.66, for details per condition See Supplementary Table 1) over all conditions. Due to experimenter error two participants were presented with only 5 blocks (1 isochronous and 1 jittered block left out, respectively). They were not rejected from the analysis, as the percentage of accepted epochs was over 50 in both conditions, and their lowest number of accepted epochs was 99 and 69 for any condition, respectively.

D1-S1 and D2-S2 difference waveforms were created for both conditions. Mean difference amplitudes were measured from the 200–300 ms time window, relative to stimulus onset (MMR). The window size and latency were determined by visual inspection.

Effects of the two conditions and two positions on MMR amplitudes were tested with a general linear model (GLM) of the structure Timing [Isochronous vs. Jittered] × Position [Beat vs. Offbeat] × Frontality [F vs. C vs. P electrode line] × Laterality [left vs. midline vs. right]. Greenhouse-Geisser correction was applied, and the corrected values together with the ε correction factor appear in the text where appropriate. η_p_^2^ effect sizes are also given in the text. The Timing × Position interaction was entered into a planned comparison. Post-hoc Tukey-HSD tests were employed to further detail other significant results. All parametric statistical analyses were computed using Statistica (version 13.4.0.14; TIBCO Software, Palo Alto, CA, U.S.A.)

In order to quantify the null result of the GLM for the Timing × Position interaction of the Jittered sequences (e.g., the absence of statistical learning in terms of the hypotheses) we used a Bayesian *t* test as implemented in JASP (JASP Team, 2019; Wagenmakers et al., 2018) to compare between the jittered beat and jittered offbeat MMR amplitudes at Cz. We estimated Bayes factors using a Cauchy prior distribution (*r* = .71) with no difference between the Beat and Off-beat MMR amplitudes as the null hypothesis. In addition, we performed a robustness check as implemented in JASP to assess whether the results would change with a different prior (*r* = 1) as originally proposed for Bayesian *t* tests (Wagenmakers et al., 2018; Jeffreys, 1961).

## Results

Figure 3 shows an overview of the central (average of C3, Cz, and C4) responses elicited by deviant and standard stimuli at the Beat and Off-beat positions in the Isochronous and Jittered conditions, together with the corresponding difference waveforms. The difference waveforms for the beat deviants appear as a broad negative wave, which is consistent with the morphology of mismatch response (MMR) in newborn infants (Kushnerenko et al., 2002). Figures including all channels included in the analyses for all grand averages (Supplementary Figures 1-3) as well as individual averages and 95% confidence intervals (CI) for all grand averages (Supplementary Figures 4-15) are given in the Supplementary Materials. Descriptive statistics are given in Supplementary Table 2. Figure 4 shows the distributions of the MMR amplitudes obtained for the Beat and Off-beat positions, separately for the Isochronous and the Jittered conditions.

**Figure 3.**
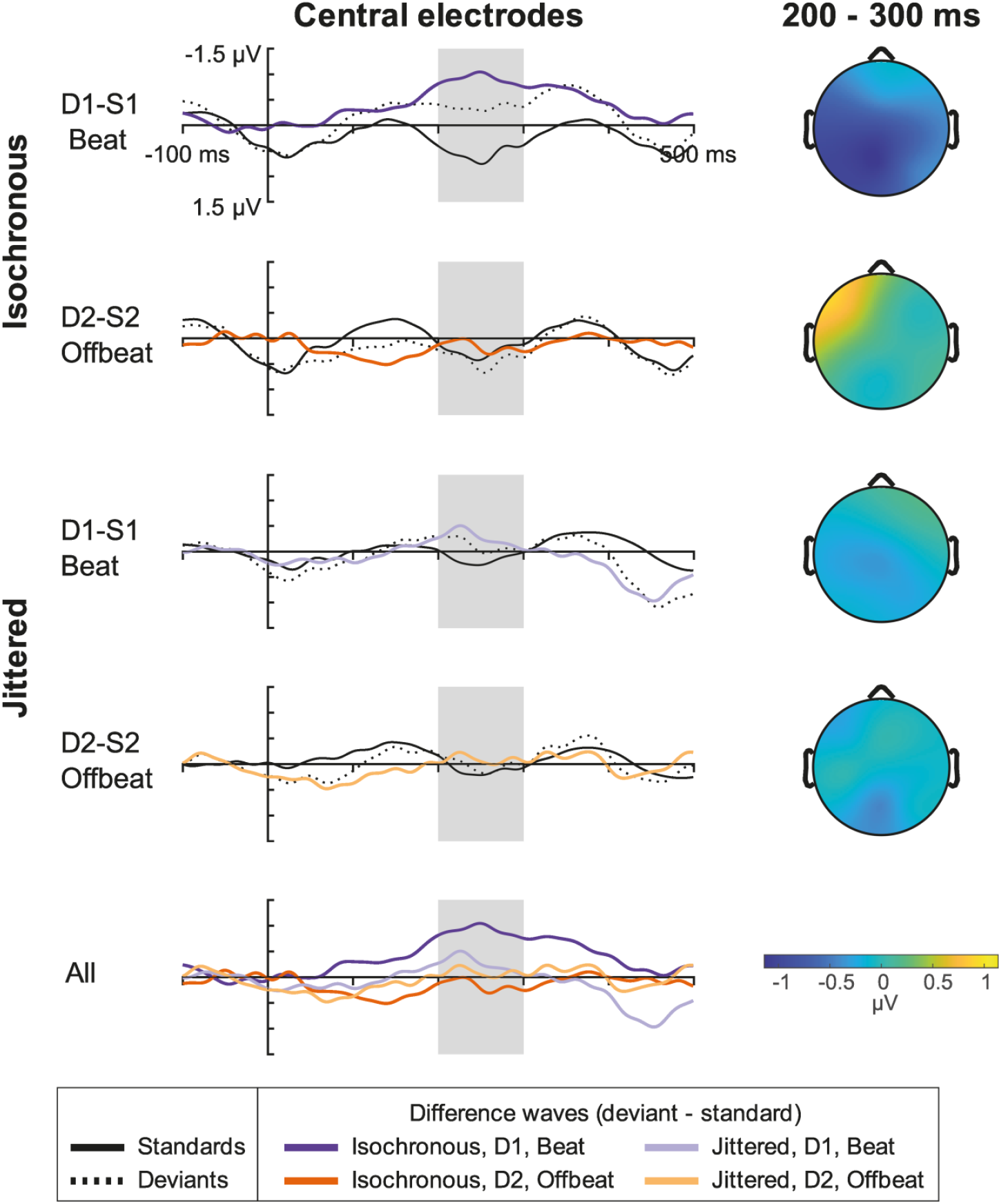
(color) Left: Grand average (N=27) ERP responses averaged over the three central electrodes (C3, Cz, C4) to deviant (D1, D2), standard (S1, S2) stimuli and the deviant minus standard difference waveforms (D1-S1, D2-S2) for the isochronous and the jittered condition. The panel on the bottom overplots all four difference waveforms. Right: Scalp distributions the average deviant-minus-standard difference amplitudes from the 200-300 ms time window (marked by grey rectangles on the signals on the left). 0 ms denotes the start of the sound at the Beat position for D1 and S1 and the start of the sound at the Offbeat position for D2 and S2.

**Figure 4.**
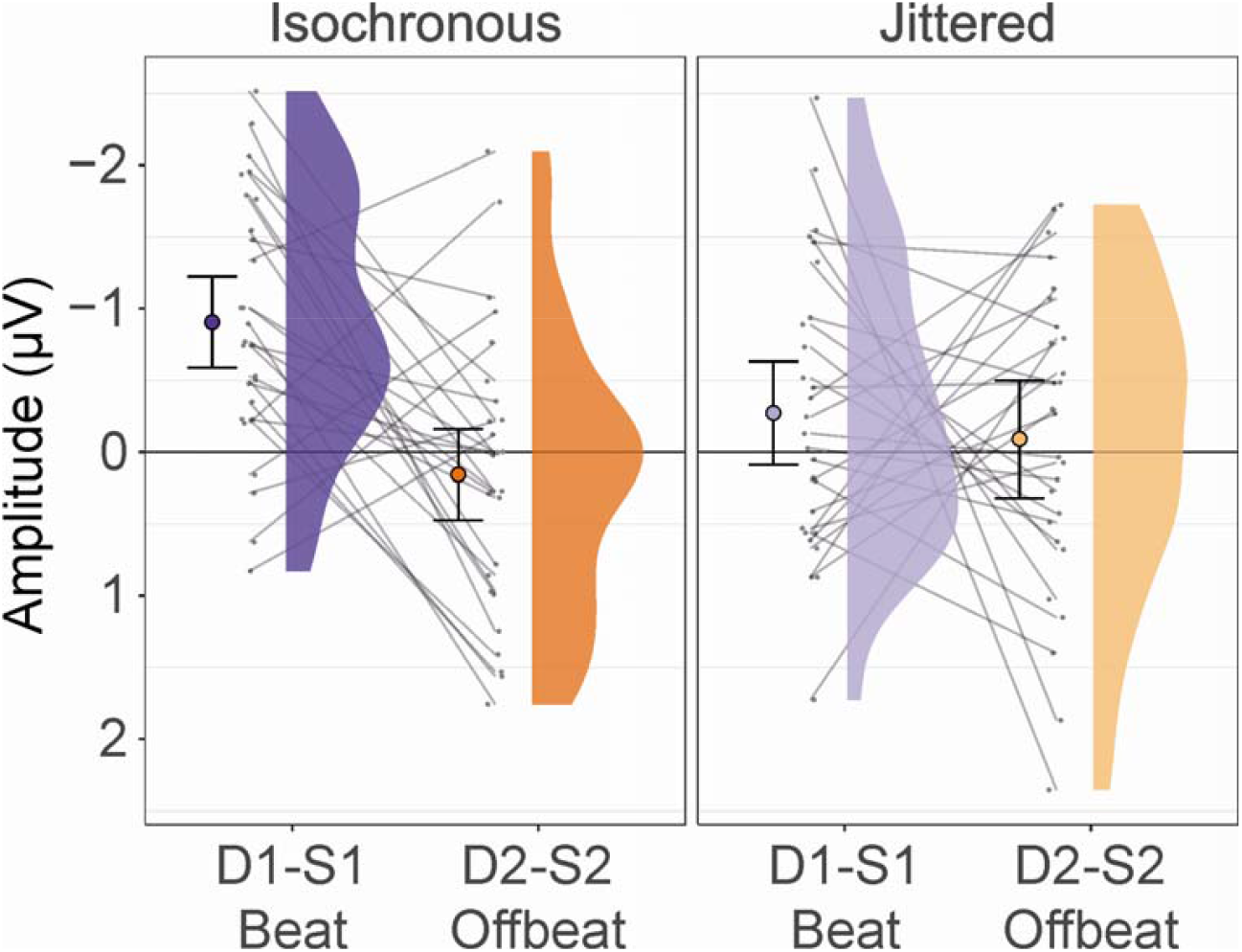
(color) Distribution of the average MMR amplitudes for all participants and conditions for the average of the three central electrodes (C3, Cz, and C4). Colored dots indicate group average with the error bars denoting 95% confidence intervals. Grey dots connected by lines show the same individual’s Beat and Off-beat MMR amplitudes, separately for the two conditions.

The Timing × Position × Frontality × Laterality model yielded a significant main effect of Position (F(1,26)= 5.086, p=0.033, η_p_^2^=0.164), which was caused by more negative MMR amplitudes for Beat compared to Offbeat positions (df=26, p=0.033), and a significant main effect of Frontality (F(1.620, 42.116)= 7.178, p= 0.004, ε= 0.810, η_p_^2^= 0.216), caused by less negative MMR amplitudes for frontal compared to both central (df=52, p=0.034) and parietal (df=52, p=0.002) electrodes.

Importantly, there was also a significant Timing × Position interaction (F(1, 26)= 5.443, p= 0.028, η_p_^2^= 0.173). Planned comparisons revealed that MMN amplitudes were significantly more negative for Beat responses, compared to Offbeat responses in the Isochronous Timing condition (F(1)= 13.136, p= 0.001) whereas they did not differ from each other in the Jittered Timing condition (F(1)= 0.013, p= 0.909) (see Figure 4). Further, Beat MMR responses were more negative in the Isochronous compared to the Jittered Timing condition (F(1)= 6.466, p= 0.017), whereas Offbeat responses did not differ between the two conditions (F(1)= 1.860, p= 0.184). The Bayesian t-test indicated moderate evidence in favor of the null hypothesis (no difference between Beat and Offbeat MMR amplitudes) for the Jittered condition (BF_01_ = 4.49). Bayes factors between 3 and 10 are regarded as moderate evidence in favor of the null hypothesis (Wagenmakers et al., 2018). The robustness check indicated that the results did not change as a function of the prior used (with the more traditional prior of *r* = 1: BF_01_ = 6.135). The output of the JASP software is included in the Supplementary Materials There was also a significant Position × Laterality effect (F(1.845, 47.962)= 3.74507576, p= 0.034, ε= 0.922, η_p_^2^= 0.126). Post-hoc testing revealed that while MMR amplitudes at the Beat position were significantly more negative compared to the Offbeat position at all electrodes, the difference was smaller between the right scalp location of the Beat position and the right and central scalp locations of the Offbeat position (df=52, p=0.044 and p= 0.034, respectively, all other differences p<0.001).

## Discussion

The results show that whereas in the isochronous condition there was a significant difference between the MMR amplitudes obtained for deviants presented at beat and offbeat positions, this difference was absent in the jittered condition. The dependence of difference responses on metrical position in the isochronous, but not the jittered condition is indicative of the presence of beat processing. These results support and provide converging evidence to the main conclusion of Winkler and colleagues (2009) that beat detection is already functional at birth in healthy infants. Importantly, the current results show that beat detection in newborns is unlikely to be solely explained by statistical learning, i.e., at least not by the known neonatal ability of learning transitional probabilities of stimulus sequences.

Beat processing (or beat perception) has been argued to be a fundamental component of the capacity for music and a prerequisite for being able to dance and make music together (Honing et al., 2015). Beat processing has been shown to be absent in macaque monkeys when using the same paradigm as in the current study (Honing et al., 2018). Beat perception (and synchronization) appears to have evolved gradually within the primates (Merchant & Honing, 2014), peaking in humans and present only with limitations in chimpanzees (Hattori & Tomonaga, 2019), bonobos (Large & Gray, 2015), and other nonhuman primates (Honing et al., 2012, Honing et al., 2018, Bouwer et al., 2021).

When combining the current results with our previous study, we now have converging evidence from two different paradigms suggesting that beat processing is functional in newborn infants (Winkler et al., 2009). This can be taken as support for a biological basis of beat perception, *per se* (Merchant & Honing, 2014; ten Cate & Honing, in press). That is, while statistical learning is an essential function for extracting information from the world, the current results suggest that it is complemented by functions honed to extract the temporal structure of (at least) the acoustic environment. Their complementary nature is supported by findings showing that temporal predictability, which is enhanced by isochronous stimulus presentation, appears to improve statistical learning (Tsogli, Jentschke & Koelsch, 2022, Selchenkova, Jones & Tillmann, 2014). In fact, much of the research on statistical learning, especially in neonates, has been based on isochronous stimuli (Bosseler et al., 2016; Teinonen et al., 2009). Indeed, it is easy to speculate that extracting the temporal structure of a sound sequence helps to reduce its variability by allowing the brain to focus its processing efforts in time (entrainment, see e.g., Stefanics et al., 2010). A similar assumption has been suggested in the Dynamic Attention Theory (Jones, 1976).

While the main hypothesis of the current study was confirmed by the data, contrary to the previous results in adults (Bouwer et al., 2016), we found no main effect of the temporal schedule of stimulus presentation (isochronous vs. jittered sound delivery). This was due to the lack of difference between offbeat MMR amplitudes in the isochronous and jittered conditions. Temporal predictability, as is present in the isochronous sequences, be it based on its regularity or on the predictable single interval (Bouwer, Nityananda, Rouse & ten Cate, 2021), should enhance processing of events in the isochronous sequence as compared to the jittered sequence (Schwartze et al., 2011). It is unlikely that newborns would be capable of detecting a beat, but not isochrony, which, in effect, is a single level of regularity. Indeed, newborns are sensitive to temporal aspects of sequences (Háden, Honing, Török, & Winkler, 2015). More likely, the absence of a difference in the offbeat positions between isochronous and jittered conditions is in fact due to beat perception. Temporal predictability in general should make events in the isochronous condition more expected (better specified), and thus violations of their regularities should elicit larger prediction errors regardless of metrical position. However, one of the main features of beat perception is that on top of enhancing the processing of metrically important events, it also suppresses processing of metrically less important ones (Bouwer et al., 2020; Breska & Deouell, 2017). Possibly, the presence of the beat in the isochronous condition of the current study hampered the processing of the offbeat events, which, in neonates, resulted in no or less specific prediction for offbeat events.

Perhaps the most striking difference between the current study in neonates and previous results with the same paradigm in adults is that in the absence of a clear MMR amplitude difference between beat and offbeat positions in the jittered condition, the current results are inconclusive with regards to the exact role of statistical learning in processing of rhythmic sequences. In the same paradigm, adults have shown different EEG responses to deviants based on position in the jittered sequences even when not attending to the stimuli (Bouwer et al., 2016). This was argued to reflect the adults’ ability to extract statistical regularity from the sequence, even when the timing is irregular. Further, statistical learning has been shown to operate in neonates (Bulf, Johnson & Valenza, 2011; Bosseler et al., 2016, Teinonen, et al., 2009). There are several possible explanations of why statistical learning was not observed in the current study. The simplest one is that our measure was not sufficiently sensitive for neonates, whose EEG has a much lower signal-to-noise ratio for ERPs than adults (Kushnerenko, Van den Bergh & Winkler, 2013). The Bayesian analysis of this null result signified only moderate evidence for no statistical learning, which leaves this question open for future investigations.

As the current data does not shed further light on this issue, here we only bring up the explanation most compatible with the notion of complementary parallel processes detecting temporal and sequential regularities in the human brain. It is possible that isochrony or, more generally, temporal predictability is important for newborns to extract sequential probabilities necessary for statistical learning. Newborns have no problem extracting even higher order regularities in temporally highly predictable stimuli (e.g., Ruusuvirta et al., 2004; Stefanics et al., 2009), and although predictability in time is not necessary for statistical learning in adults, it does enhance the statistical MMN amplitude, an index of statistical learning (Tsogli, Jentschke & Koelsch, 2022). Further, in general, the noisier the regularity, the lower the MMN amplitude (e.g., Winkler et al., 1990). In other words, predictions with low specificity result in low-amplitude error signals (Friston, 2005; Southwell & Chait, 2018). If, as we have hypothesized above, extracting temporal structure reduces the observed variability (e.g., by separating the regularities for beat and offbeat sounds) while helping to focus the processing in time (entrainment), it may substantially reduce the necessary capacities, which may be even more crucial for neonates than for adults. Obviously, further studies are needed to test this hypothesis.

Some indirect support for this hypothesis may be gained from the observed significant Position × Laterality interaction. The processing of different auditory features is thought to be lateralized (e.g., Tervaniemi & Hugdahl, 2003): left for temporal features and right for spectral features (Zatorre & Belin, 2001), which seems to be the case also for newborns (DeCasper & Prescott, 2009). Entrainment to a structured sequence in a statistical learning task showed similar lateralization of brain activity to the current one compared to a non-structured sequence in adults with better performance (Moser et al., 2021). This brings up the possibility of an interaction between entrainment and statistical learning, as we have proposed above.

## Conclusions

In the current experiment we have established that newborn infants are capable of beat based processing, providing converging evidence for the conclusions of Winkler et al. (2009). Importantly, the paradigm of Bouwer et al. (2016) used here allowed for the separation of beat processing and statistical learning of transition probabilities in neonates. Although the results suggest the presence of beat perception in newborns, we could not show the presence of statistical learning of transition probabilities when sequence timing was irregular. Current results, and previous results that show better statistical learning for sequences of regular temporal structure, bring up the possibility that extracting the temporal structure and statistical learning work in a complementary fashion.

## Supporting information

Sample sound 1

Sample sound 2

## Acknowledgement

The authors are grateful to all infants and their parents for participating in the experiments, Prof. Dr. Miklós Török and Dr József Korcsik for supporting the experiments at the Department of Obstetrics, Military Hospital, Budapest, and Judit Roschéné Farkas for recording the data.

