## Additional tables and figures for "Beat processing in newborn infants cannot be explained by statistical learning based on transition probabilities"

Supplementary Materials

**Supplementary Table 1**

Presented and accepted number of epochs per condition, their percentage with mean and standard deviation

| Condition | D1 (Beat Deviant Isochronous) | | | D2 (Offbeat Deviant Isochronous) | | | D1 (Beat Deviant Jittered) | | | D2 (Offbeat Deviant Jittered) | | |
| --- | --- | --- | --- | --- | --- | --- | --- | --- | --- | --- | --- | --- |
| N Epochs | Presented | Accepted | Percentage | Presented | Accepted | Percentage | Presented | Accepted | Percentage | Presented | Accepted | Percentage |
| Participant |  |  |  |  |  |  |  |  |  |  |  |  |
| 1585 | 195 | 166 | 85.13 | 195 | 172 | 88.21 | 195 | 168 | 86.15 | 195 | 177 | 90.77 |
| 1586 | 195 | 98 | 50.26 | 195 | 100 | 51.28 | 195 | 134 | 68.72 | 195 | 149 | 76.41 |
| 1588 | 195 | 103 | 52.82 | 195 | 105 | 53.85 | 195 | 144 | 73.85 | 195 | 133 | 68.21 |
| 1589 | 195 | 147 | 75.38 | 195 | 158 | 81.03 | 130 | 69 | 53.08 | 130 | 73 | 56.15 |
| 1590 | 195 | 142 | 72.82 | 194 | 144 | 74.23 | 195 | 134 | 68.72 | 195 | 151 | 77.44 |
| 1593 | 195 | 122 | 62.56 | 195 | 131 | 67.18 | 195 | 107 | 54.87 | 195 | 104 | 53.33 |
| 1594 | 195 | 135 | 69.23 | 195 | 131 | 67.18 | 195 | 141 | 72.31 | 195 | 145 | 74.36 |
| 1595 | 195 | 164 | 84.10 | 195 | 153 | 78.46 | 195 | 166 | 85.13 | 195 | 166 | 85.13 |
| 1596 | 195 | 187 | 95.90 | 194 | 185 | 95.36 | 195 | 141 | 72.31 | 195 | 131 | 67.18 |
| 1598 | 195 | 126 | 64.62 | 195 | 141 | 72.31 | 195 | 165 | 84.62 | 195 | 167 | 85.64 |
| 1600 | 195 | 174 | 89.23 | 195 | 168 | 86.15 | 195 | 129 | 66.15 | 194 | 130 | 67.01 |
| 1602 | 195 | 175 | 89.74 | 195 | 173 | 88.72 | 195 | 151 | 77.44 | 195 | 159 | 81.54 |
| 1604 | 195 | 188 | 96.41 | 195 | 193 | 98.97 | 195 | 183 | 93.85 | 195 | 180 | 92.31 |
| 1607 | 195 | 188 | 96.41 | 195 | 184 | 94.36 | 195 | 164 | 84.10 | 195 | 170 | 87.18 |
| 1608 | 195 | 192 | 98.46 | 195 | 191 | 97.95 | 195 | 190 | 97.44 | 195 | 189 | 96.92 |
| 1609 | 195 | 187 | 95.90 | 195 | 180 | 92.31 | 195 | 186 | 95.38 | 195 | 193 | 98.97 |
| 1610 | 195 | 181 | 92.82 | 195 | 174 | 89.23 | 195 | 188 | 96.41 | 195 | 182 | 93.33 |
| 1612 | 130 | 109 | 83.85 | 130 | 99 | 76.15 | 195 | 172 | 88.21 | 195 | 167 | 85.64 |
| 1613 | 195 | 165 | 84.62 | 195 | 170 | 87.18 | 195 | 171 | 87.69 | 195 | 170 | 87.18 |
| 1615 | 195 | 187 | 95.90 | 195 | 182 | 93.33 | 195 | 195 | 100.00 | 195 | 195 | 100.00 |
| 1617 | 195 | 181 | 92.82 | 195 | 179 | 91.79 | 195 | 190 | 97.44 | 195 | 189 | 96.92 |
| 1618 | 195 | 160 | 82.05 | 195 | 154 | 78.97 | 195 | 151 | 77.44 | 195 | 146 | 74.87 |
| 1619 | 195 | 191 | 97.95 | 195 | 188 | 96.41 | 195 | 192 | 98.46 | 195 | 191 | 97.95 |
| 1621 | 195 | 161 | 82.56 | 195 | 165 | 84.62 | 195 | 160 | 82.05 | 195 | 172 | 88.21 |
| 1622 | 195 | 171 | 87.69 | 195 | 177 | 90.77 | 195 | 189 | 96.92 | 195 | 191 | 97.95 |
| 1623 | 195 | 134 | 68.72 | 195 | 146 | 74.87 | 195 | 139 | 71.28 | 195 | 157 | 80.51 |
| 1625 | 195 | 189 | 96.92 | 195 | 191 | 97.95 | 195 | 176 | 90.26 | 195 | 169 | 86.67 |
| Mean | 193 | 160 | 83.1 | 193 | 161 | 83.3 | 193 | 159 | 82.23 | 193 | 161 | 83.25 |
| SD | 12 | 29 | 13.8 | 12 | 27 | 12.7 | 12 | 29 | 13.03 | 12 | 28 | 12.58 |

**Supplementary Table 1 (continued)**

| Condition | D1 (Beat Standard Isochronous) | | | S2 (Offbeat Standard Isochronous) | | | S1 (Beat Standard Jittered) | | | S2 (Offbeat Standard Jittered) | | |
| --- | --- | --- | --- | --- | --- | --- | --- | --- | --- | --- | --- | --- |
| N Epochs | Presented | Accepted | Percentage | Presented | Accepted | Percentage | Presented | Accepted | Percentage | Presented | Accepted | Percentage |
| Participant |  |  |  |  |  |  |  |  |  |  |  |  |
| 1585 | 975 | 843 | 86.46 | 845 | 728 | 86.15 | 975 | 855 | 87.69 | 845 | 742 | 87.81 |
| 1586 | 984 | 503 | 51.12 | 843 | 432 | 51.25 | 984 | 688 | 69.92 | 843 | 598 | 70.94 |
| 1588 | 983 | 554 | 56.36 | 844 | 478 | 56.64 | 983 | 706 | 71.82 | 844 | 594 | 70.38 |
| 1589 | 976 | 767 | 78.59 | 845 | 666 | 78.82 | 653 | 356 | 54.52 | 564 | 310 | 54.96 |
| 1590 | 975 | 696 | 71.38 | 840 | 606 | 72.14 | 979 | 687 | 70.17 | 845 | 612 | 72.43 |
| 1593 | 976 | 645 | 66.09 | 846 | 557 | 65.84 | 976 | 549 | 56.25 | 846 | 473 | 55.91 |
| 1594 | 976 | 691 | 70.80 | 844 | 597 | 70.73 | 976 | 718 | 73.57 | 844 | 628 | 74.41 |
| 1595 | 970 | 780 | 80.41 | 843 | 673 | 79.83 | 970 | 809 | 83.40 | 843 | 713 | 84.58 |
| 1596 | 967 | 937 | 96.90 | 841 | 819 | 97.38 | 968 | 667 | 68.90 | 842 | 565 | 67.10 |
| 1598 | 976 | 720 | 73.77 | 845 | 634 | 75.03 | 976 | 849 | 86.99 | 845 | 725 | 85.80 |
| 1600 | 974 | 844 | 86.65 | 846 | 724 | 85.58 | 973 | 686 | 70.50 | 845 | 611 | 72.31 |
| 1602 | 976 | 885 | 90.68 | 842 | 767 | 91.09 | 976 | 794 | 81.35 | 842 | 681 | 80.88 |
| 1604 | 967 | 942 | 97.41 | 845 | 816 | 96.57 | 967 | 891 | 92.14 | 845 | 767 | 90.77 |
| 1607 | 976 | 919 | 94.16 | 844 | 784 | 92.89 | 976 | 849 | 86.99 | 844 | 747 | 88.51 |
| 1608 | 991 | 978 | 98.69 | 845 | 831 | 98.34 | 991 | 964 | 97.28 | 845 | 824 | 97.51 |
| 1609 | 970 | 911 | 93.92 | 844 | 792 | 93.84 | 970 | 930 | 95.88 | 844 | 806 | 95.50 |
| 1610 | 987 | 886 | 89.77 | 845 | 762 | 90.18 | 987 | 943 | 95.54 | 845 | 806 | 95.38 |
| 1612 | 652 | 519 | 79.60 | 562 | 447 | 79.54 | 977 | 856 | 87.62 | 844 | 727 | 86.14 |
| 1613 | 980 | 851 | 86.84 | 844 | 727 | 86.14 | 980 | 861 | 87.86 | 844 | 741 | 87.80 |
| 1615 | 979 | 929 | 94.89 | 844 | 797 | 94.43 | 979 | 967 | 98.77 | 844 | 835 | 98.93 |
| 1617 | 975 | 904 | 92.72 | 842 | 779 | 92.52 | 975 | 940 | 96.41 | 842 | 812 | 96.44 |
| 1618 | 973 | 770 | 79.14 | 842 | 664 | 78.86 | 973 | 757 | 77.80 | 842 | 647 | 76.84 |
| 1619 | 986 | 948 | 96.15 | 842 | 811 | 96.32 | 986 | 958 | 97.16 | 842 | 812 | 96.44 |
| 1621 | 994 | 832 | 83.70 | 848 | 724 | 85.38 | 994 | 834 | 83.90 | 848 | 716 | 84.43 |
| 1622 | 981 | 860 | 87.67 | 843 | 749 | 88.85 | 981 | 946 | 96.43 | 843 | 814 | 96.56 |
| 1623 | 975 | 689 | 70.67 | 845 | 596 | 70.53 | 975 | 749 | 76.82 | 845 | 660 | 78.11 |
| 1625 | 976 | 931 | 95.39 | 844 | 804 | 95.26 | 976 | 854 | 87.50 | 844 | 735 | 87.09 |
| Mean | 966 | 805 | 83.3 | 833 | 695 | 83.3 | 966 | 802 | 82.71 | 834 | 693 | 82.74 |
| SD | 62 | 134 | 12.5 | 53 | 115 | 12.3 | 62 | 139 | 12.23 | 53 | 118 | 12.07 |

**Supplementary Table 2**

Descriptive statistics for the mean difference waves in the 200-300ms time window per condition

| Beat | Isochronous | |  |  |  |
| --- | --- | --- | --- | --- | --- |
| Channel | Valid N | Mean | Minimum | Maximum | Std.Dev. |
| F3 | 27 | -0.55704 | -2.511 | 1.021 | 1.010343 |
| Fz | 27 | -0.34956 | -2.602 | 1.623 | 0.953322 |
| F4 | 27 | -0.32456 | -2.723 | 1.821 | 1.05179 |
| C3 | 27 | -1.01119 | -2.958 | 0.329 | 0.946282 |
| Cz | 27 | -1.02385 | -3.223 | 1.251 | 1.171495 |
| C4 | 27 | -0.68559 | -2.859 | 1.329 | 0.999737 |
| P3 | 27 | -1.09874 | -3.601 | 1.525 | 1.136376 |
| Pz | 27 | -1.13548 | -3.552 | 0.923 | 1.249976 |
| P4 | 27 | -0.60278 | -2.593 | 1.739 | 1.22202 |

| Offbeat | Isochronous | |  |  |  |
| --- | --- | --- | --- | --- | --- |
| Channel | Valid N | Mean | Minimum | Maximum | Std.Dev. |
| F3 | 27 | 0.743333 | -1.096 | 2.701 | 0.929959 |
| Fz | 27 | 0.252593 | -1.536 | 1.616 | 0.862689 |
| F4 | 27 | -0.02407 | -3.417 | 1.798 | 1.025803 |
| C3 | 27 | 0.3 | -1.479 | 1.851 | 0.932223 |
| Cz | 27 | 0.109926 | -2.391 | 3.008 | 1.526452 |
| C4 | 27 | 0.059111 | -2.416 | 2.073 | 1.132106 |
| P3 | 27 | -0.02511 | -2.643 | 2.25 | 1.1334 |
| Pz | 27 | -0.0877 | -4.878 | 3.44 | 1.648547 |
| P4 | 27 | 0.040074 | -2.544 | 2.445 | 1.078039 |

| Beat | Jittered |  |  |  |  |
| --- | --- | --- | --- | --- | --- |
| Channel | Valid N | Mean | Minimum | Maximum | Std.Dev. |
| F3 | 27 | -0.15189 | -2.047 | 2.523 | 0.962698 |
| Fz | 27 | -0.06559 | -2.244 | 2.396 | 1.245574 |
| F4 | 27 | 0.092481 | -1.938 | 2.3 | 1.050983 |
| C3 | 27 | -0.30459 | -2.749 | 2.336 | 1.129477 |
| Cz | 27 | -0.34615 | -3.761 | 1.81 | 1.364561 |
| C4 | 27 | -0.16615 | -2.114 | 2.26 | 1.104165 |
| P3 | 27 | -0.20459 | -2.533 | 2.098 | 1.21984 |
| Pz | 27 | -0.26404 | -3.696 | 2.528 | 1.339615 |
| P4 | 27 | -0.27307 | -3.622 | 4.008 | 1.439503 |

**Supplementary Table 2 (continued)**

| Offbeat | Jittered |  |  |  |  |
| --- | --- | --- | --- | --- | --- |
| Channel | Valid N | Mean | Minimum | Maximum | Std.Dev. |
| F3 | 27 | -0.20985 | -2.495 | 1.435 | 1.032062 |
| Fz | 27 | 0.017667 | -2.606 | 2.617 | 1.239148 |
| F4 | 27 | -0.03052 | -1.602 | 2.149 | 0.909022 |
| C3 | 27 | 0.004889 | -1.889 | 2.323 | 1.043516 |
| Cz | 27 | -0.14144 | -2.966 | 3.457 | 1.626637 |
| C4 | 27 | -0.13633 | -2.532 | 1.824 | 1.065358 |
| P3 | 27 | -0.27904 | -5.563 | 2.017 | 1.514037 |
| Pz | 27 | -0.44393 | -4.53 | 2.747 | 1.446434 |
| P4 | 27 | -0.14496 | -2.94 | 3.191 | 1.456851 |

**Supplementary Figure 1**

Grand average differences waves for all conditions

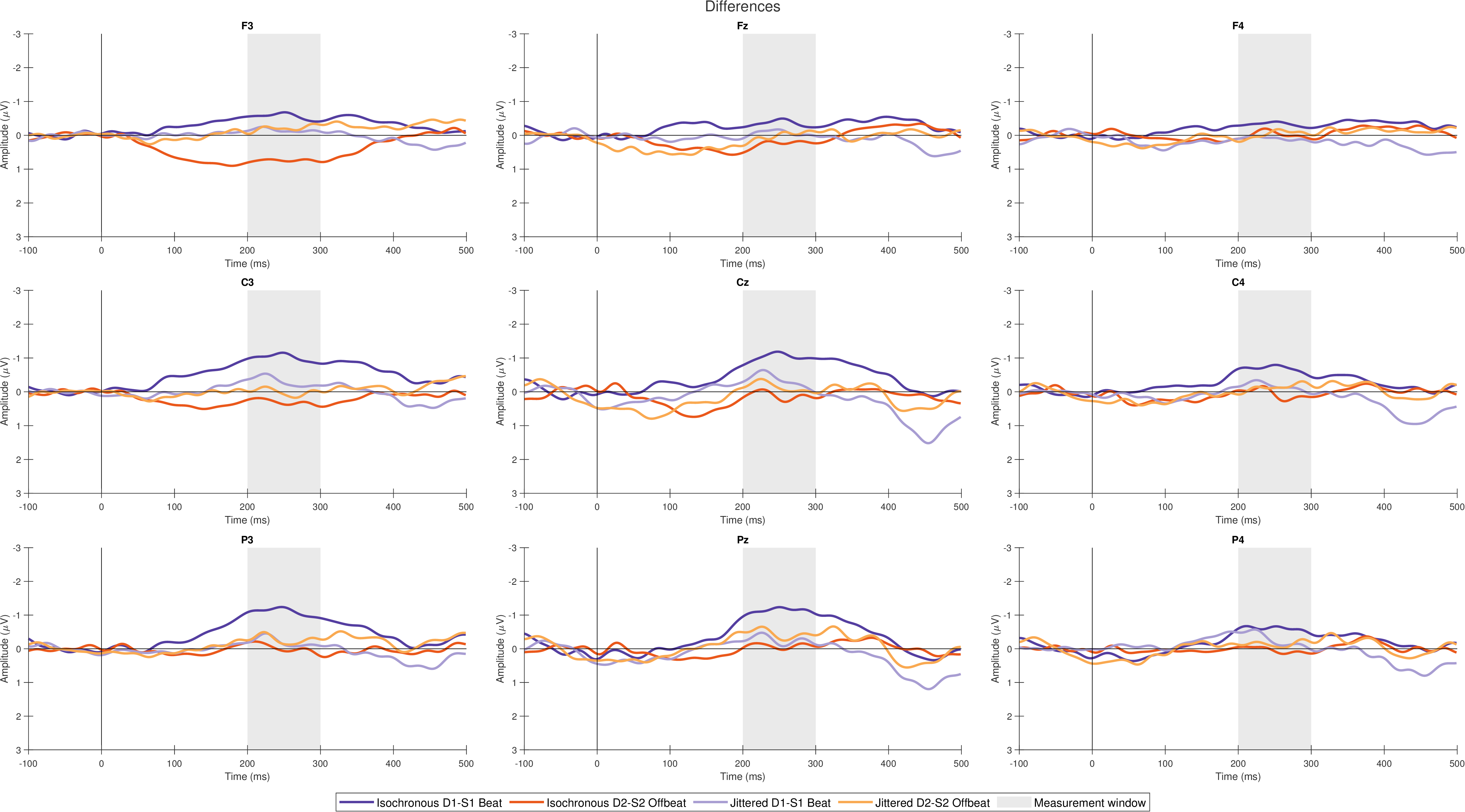

**Supplementary Figure 2**

Grand average deviant waves for all conditions

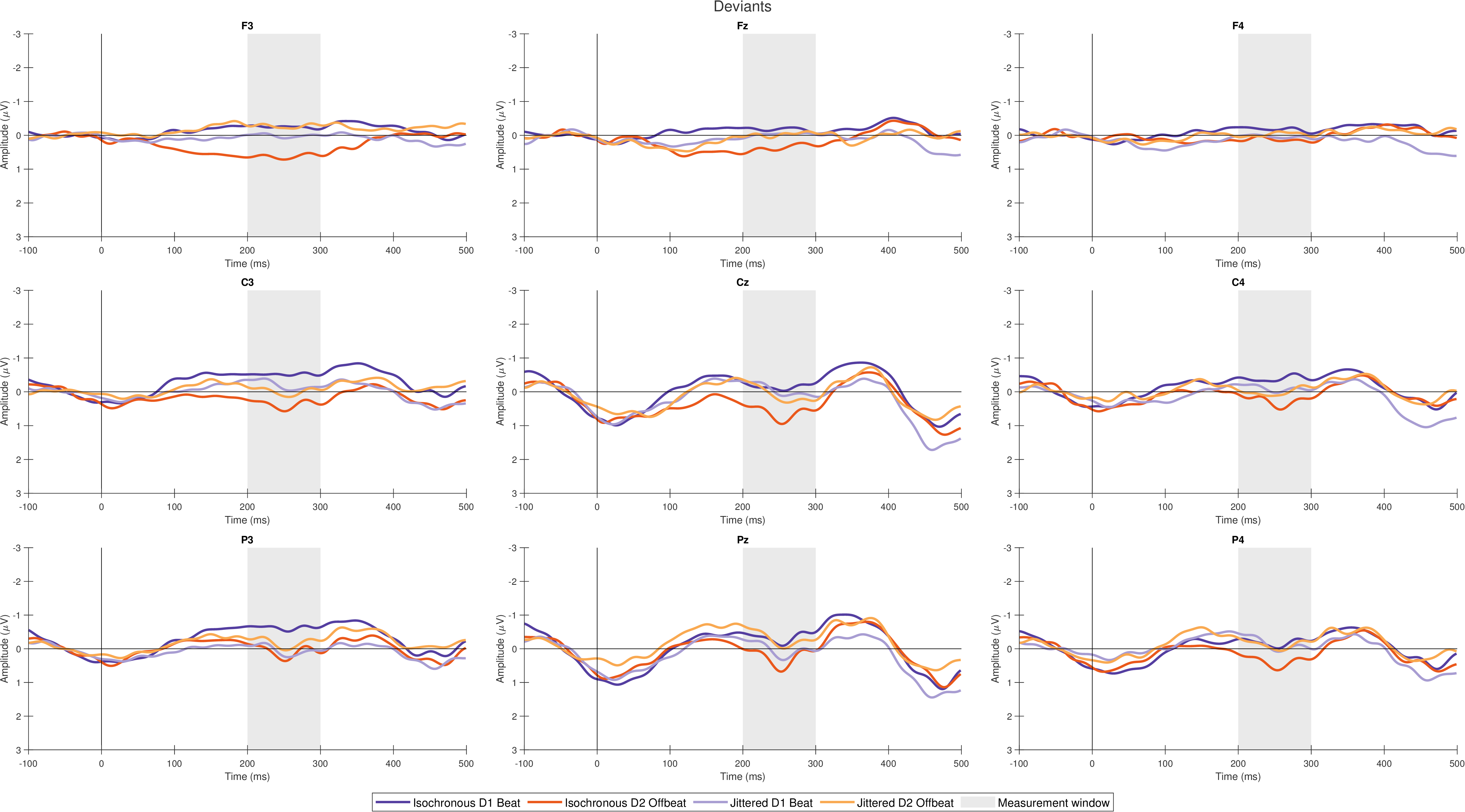

**Supplementary Figure 3**

Grand average standard waves for all conditions

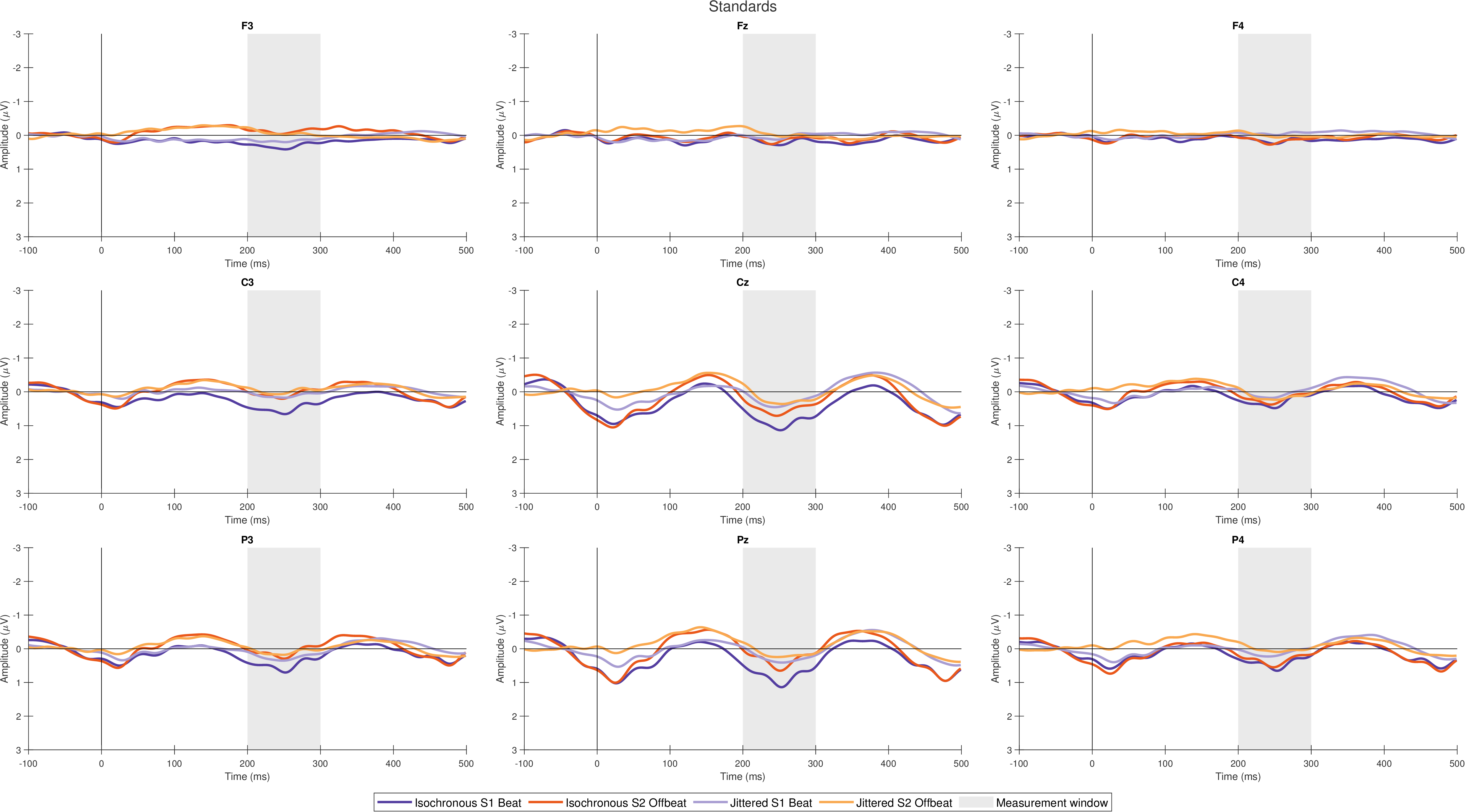

**Supplementary Figure 4**

Individual difference waves (grey) and grand average wave with 95% confidence intervals (color), Isochronous D1-S1 Beat

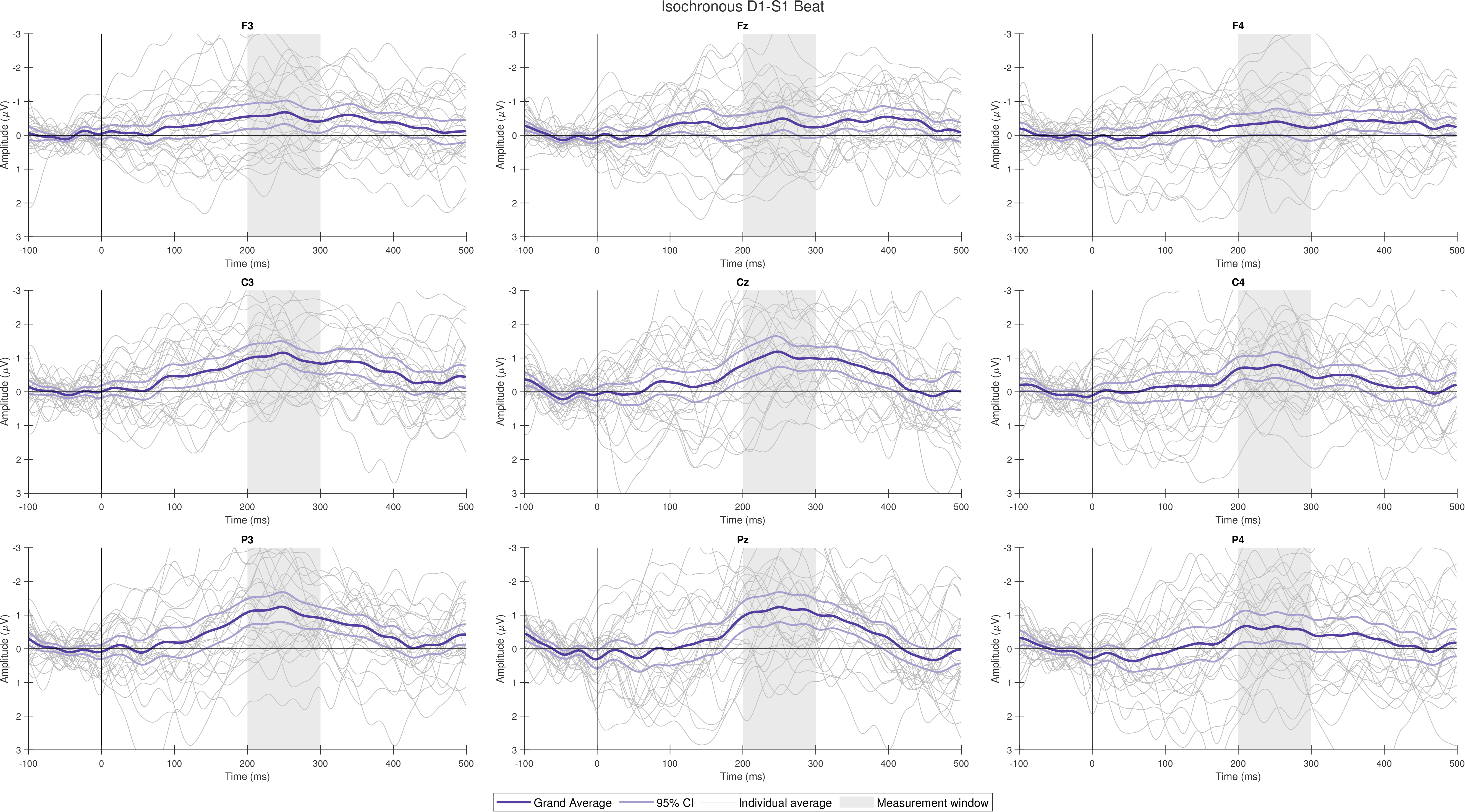

**Supplementary Figure 5**

Individual difference waves (grey) and grand average wave with 95% confidence intervals (color), Isochronous D2-S2 Offbeat

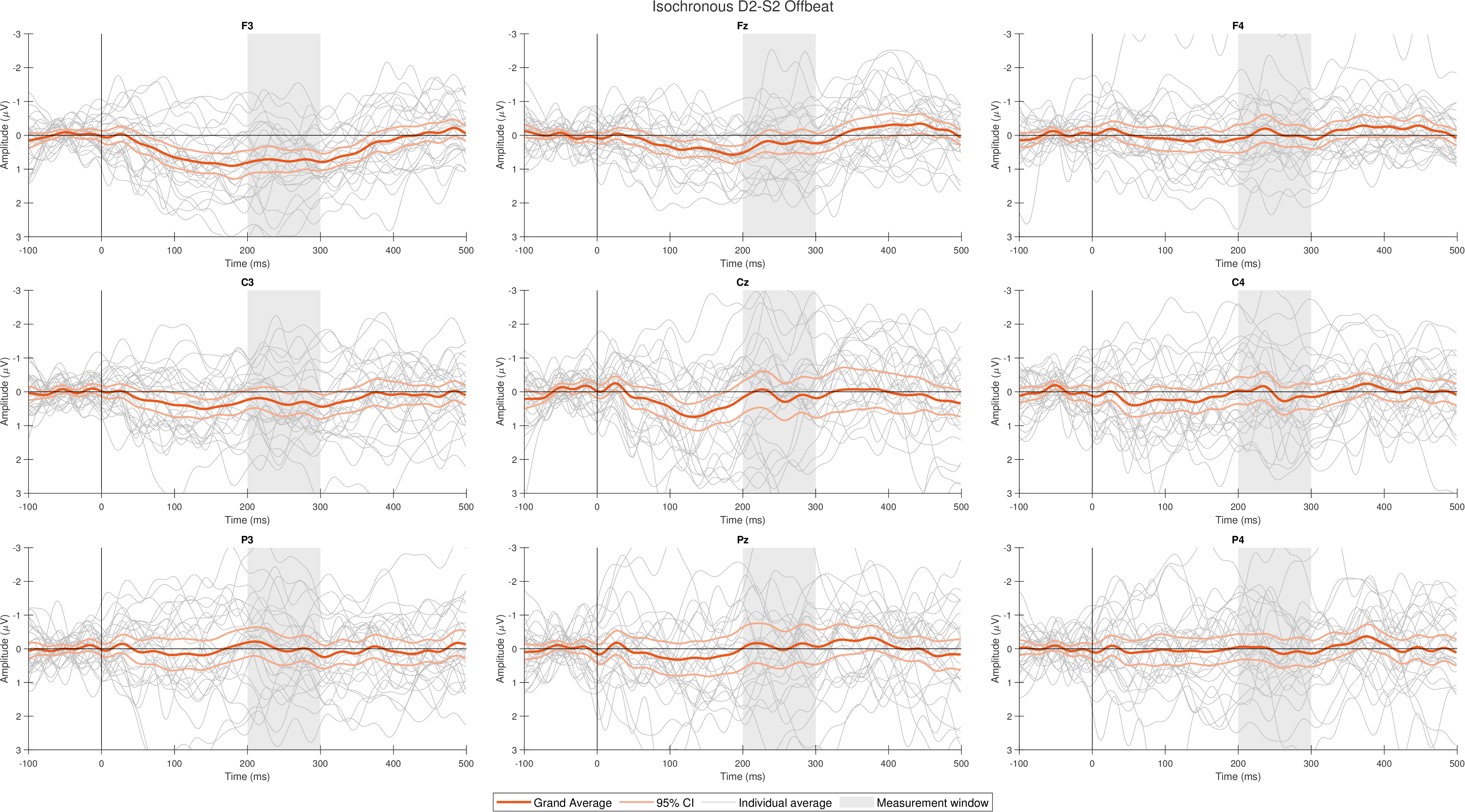

**Supplementary Figure 6**

Individual difference waves (grey) and grand average wave with 95% confidence intervals (color), Isochronous D1 Beat

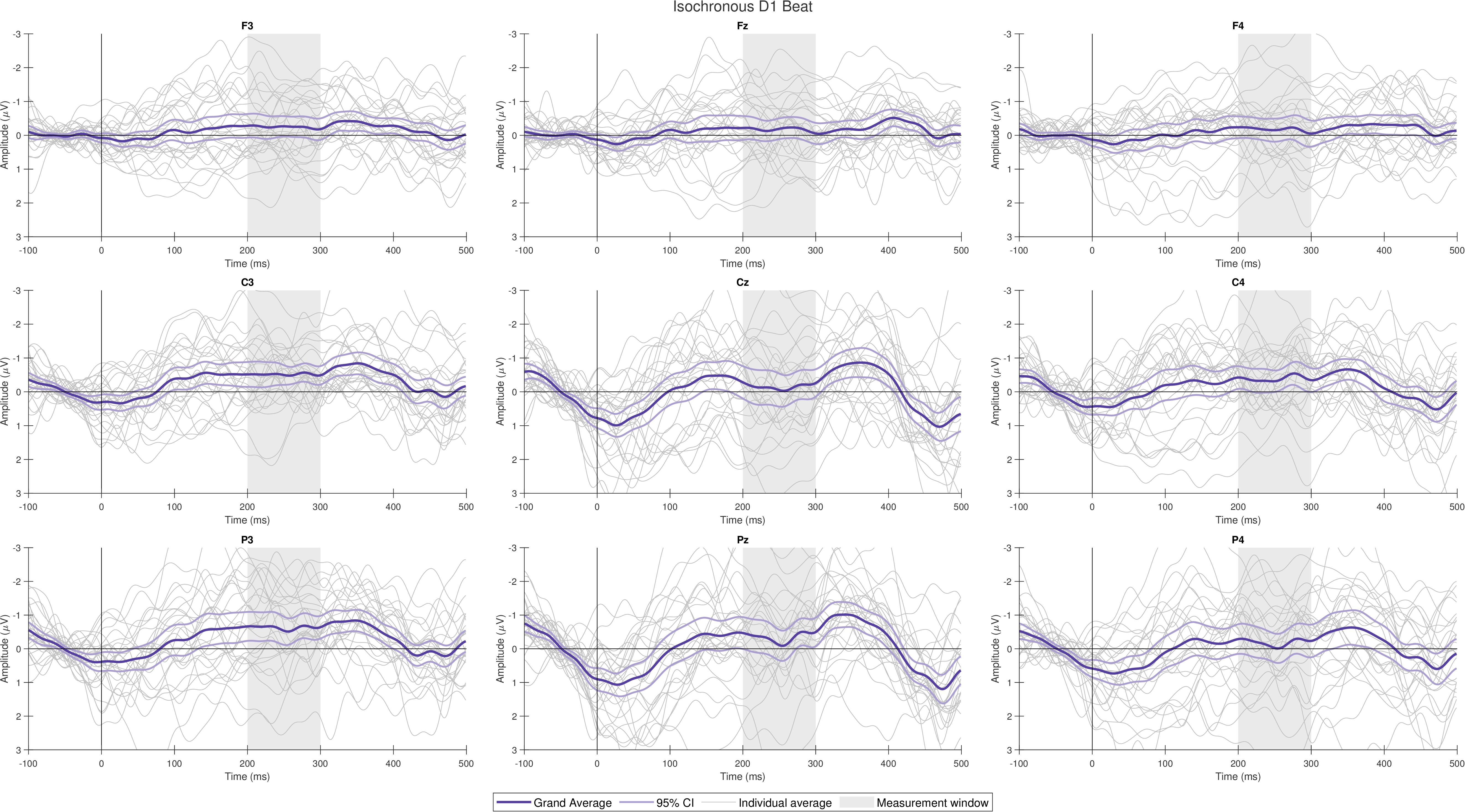

**Supplementary Figure 7**

Individual difference waves (grey) and grand average wave with 95% confidence intervals (color), Isochronous D2 Offbeat

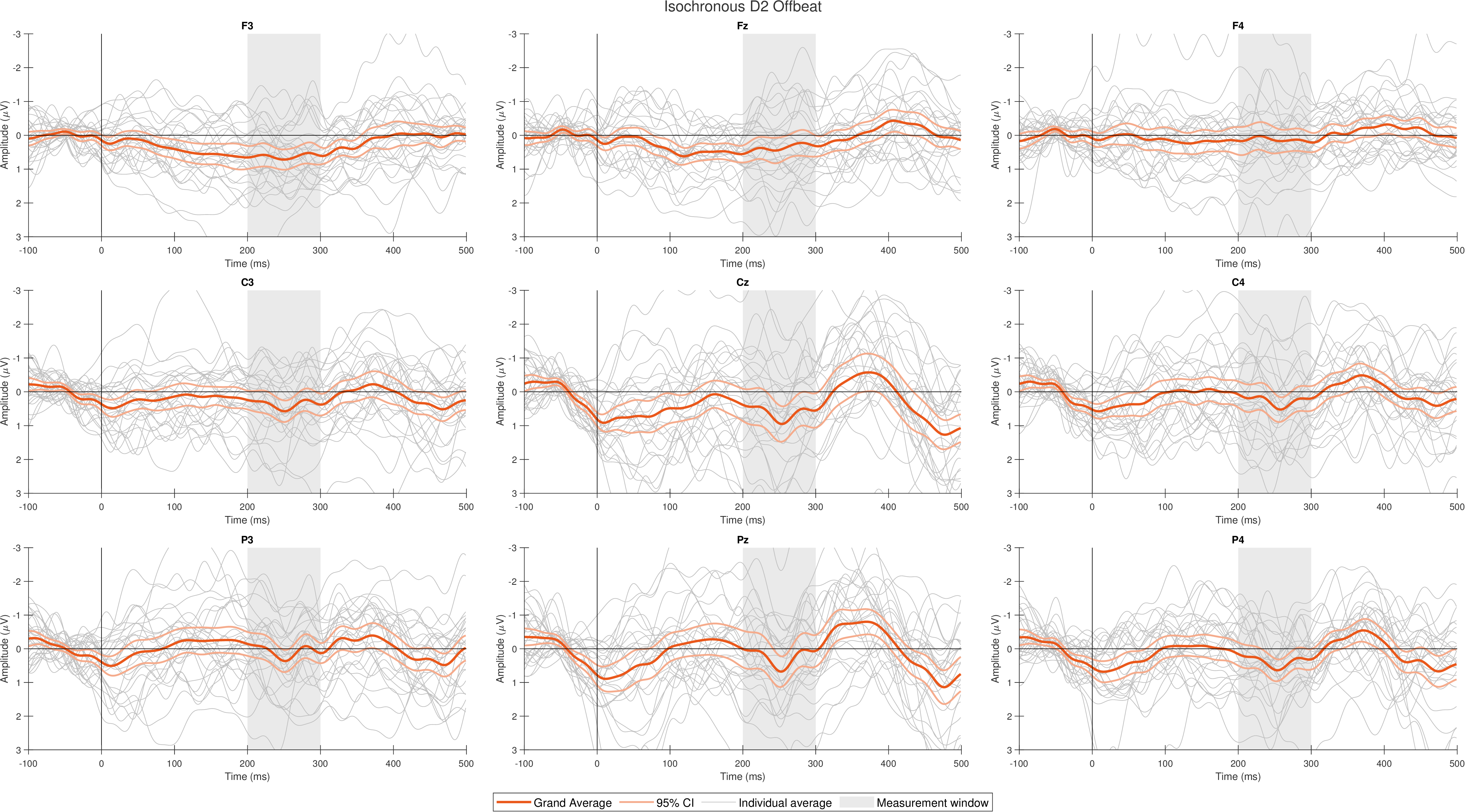

**Supplementary Figure 8**

Individual difference waves (grey) and grand average wave with 95% confidence intervals (color), Isochronous S1 Beat

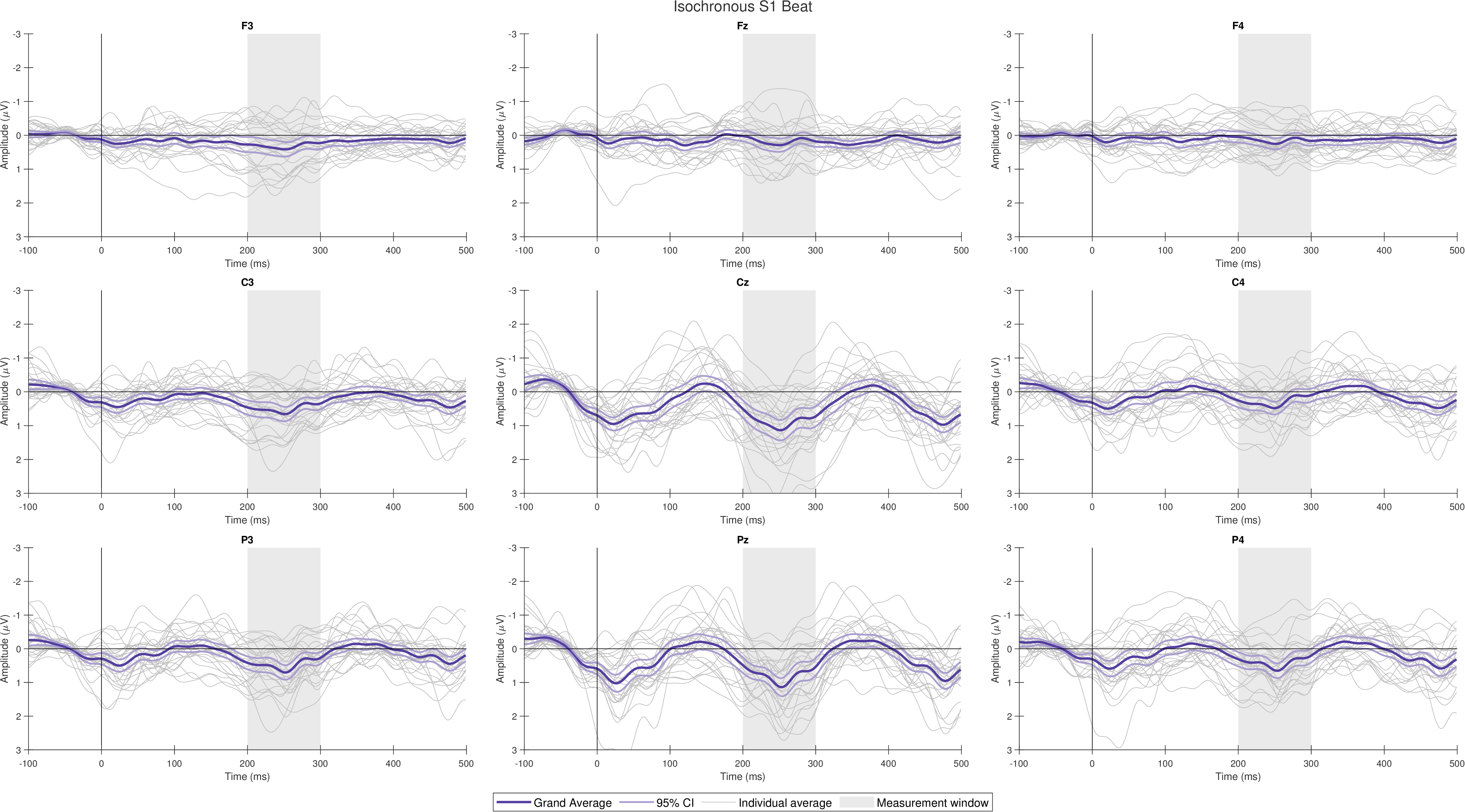

**Supplementary Figure 9**

Individual difference waves (grey) and grand average wave with 95% confidence intervals (color), Isochronous S2 Offbeat

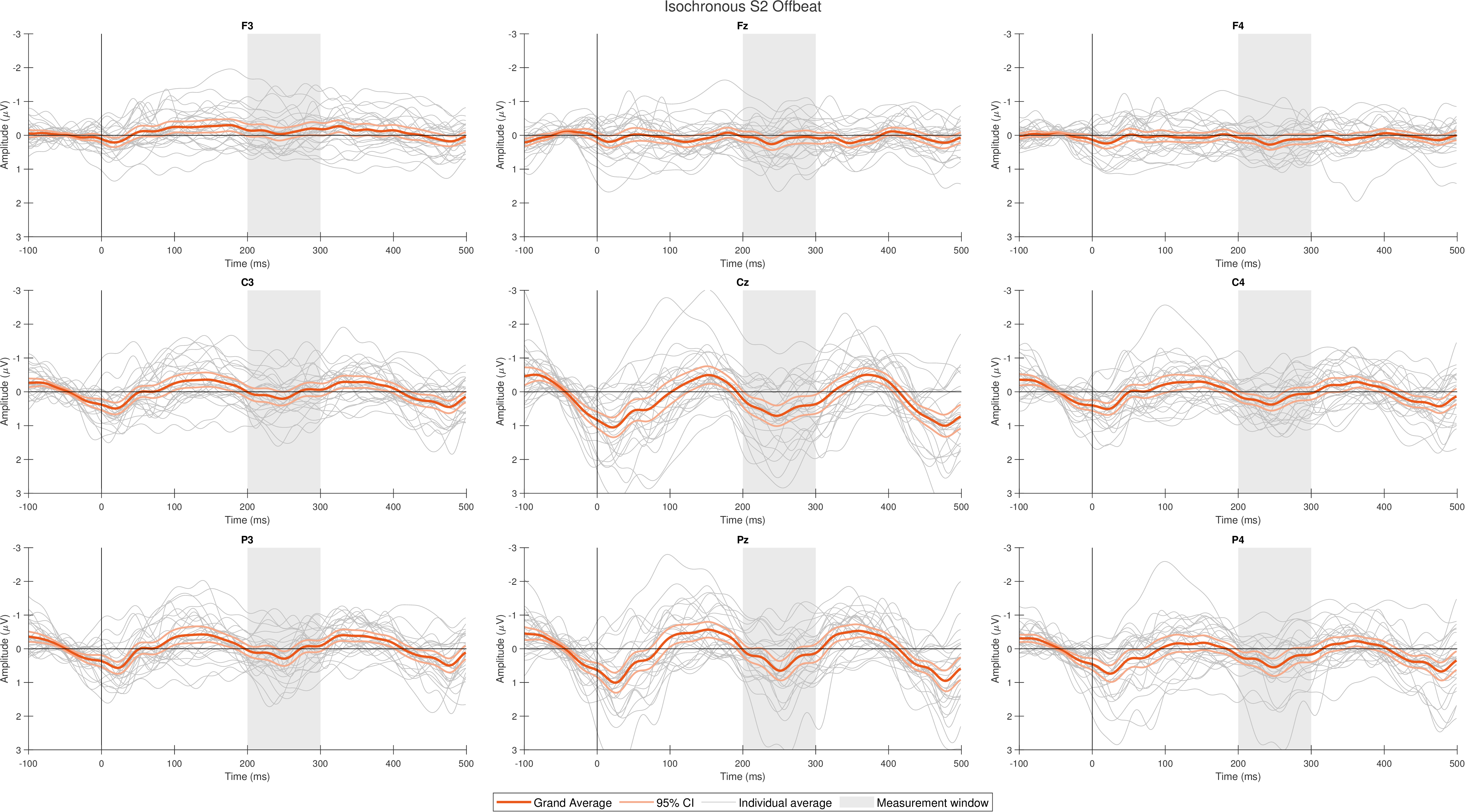

**Supplementary Figure 10**

Individual difference waves (grey) and grand average wave with 95% confidence intervals (color), Jittered D1-S1 Beat

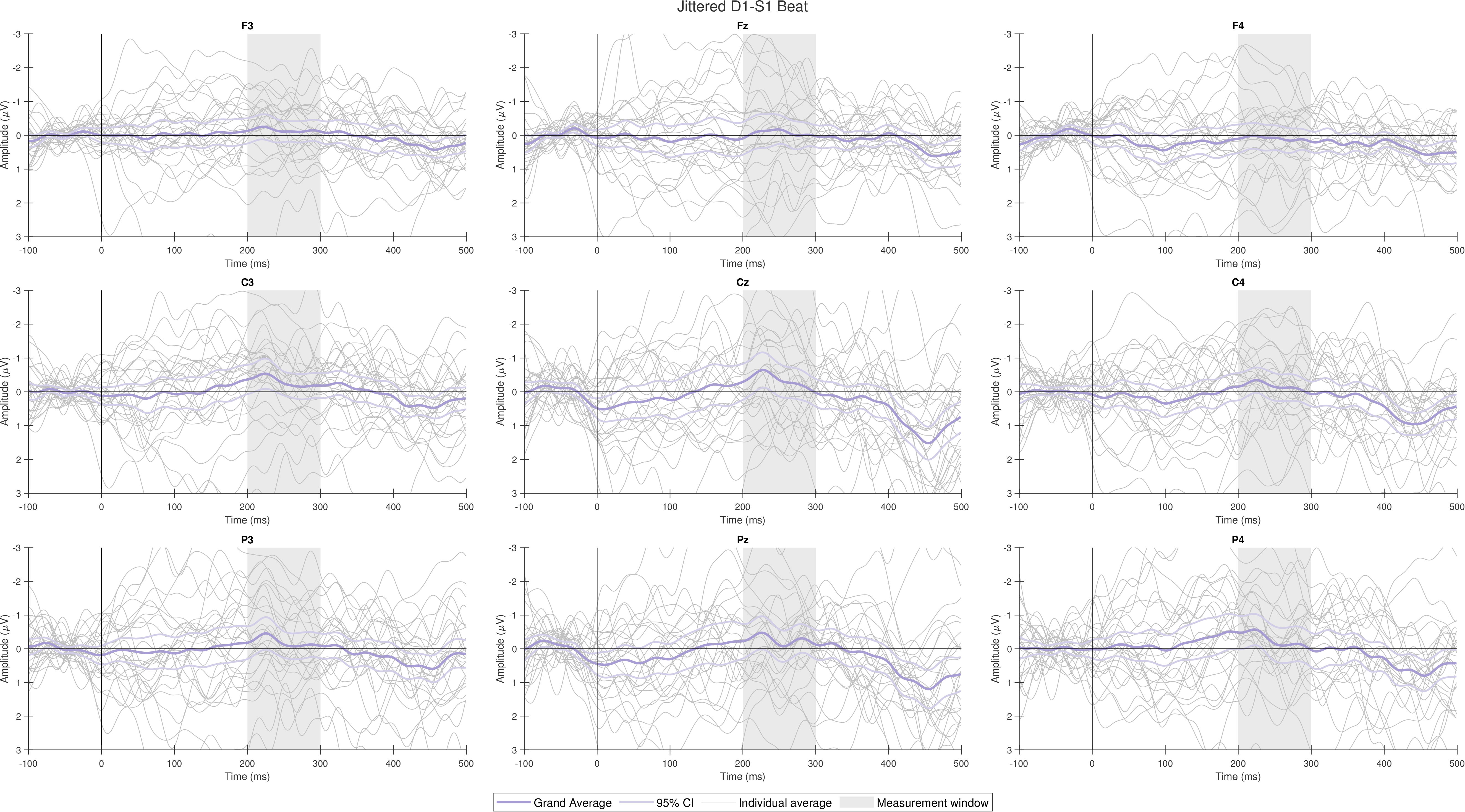

**Supplementary Figure 11**

Individual difference waves (grey) and grand average wave with 95% confidence intervals (color), Jittered D2-S2 Offbeat

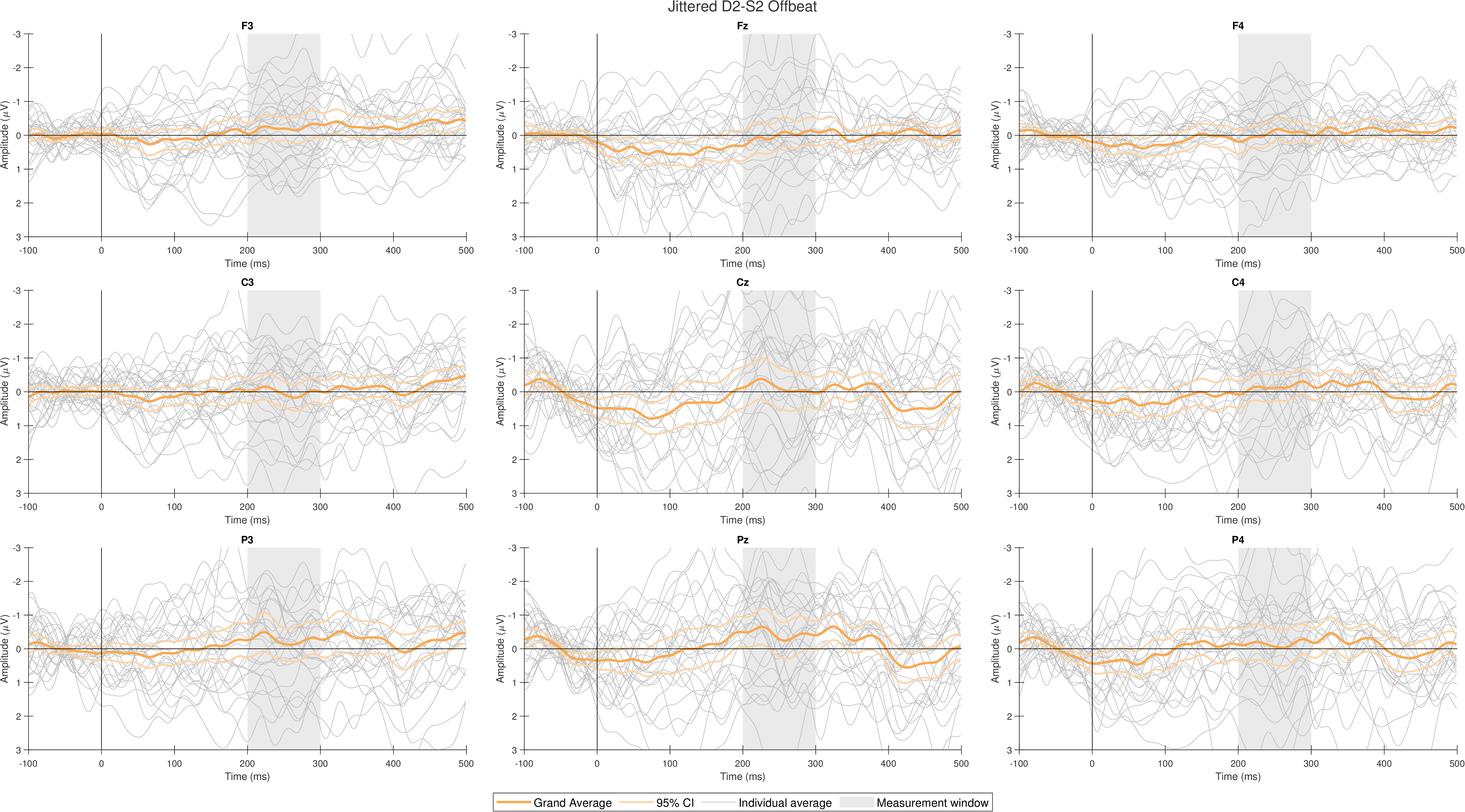

**Supplementary Figure 12**

Individual difference waves (grey) and grand average wave with 95% confidence intervals (color), Jittered D1 Beat

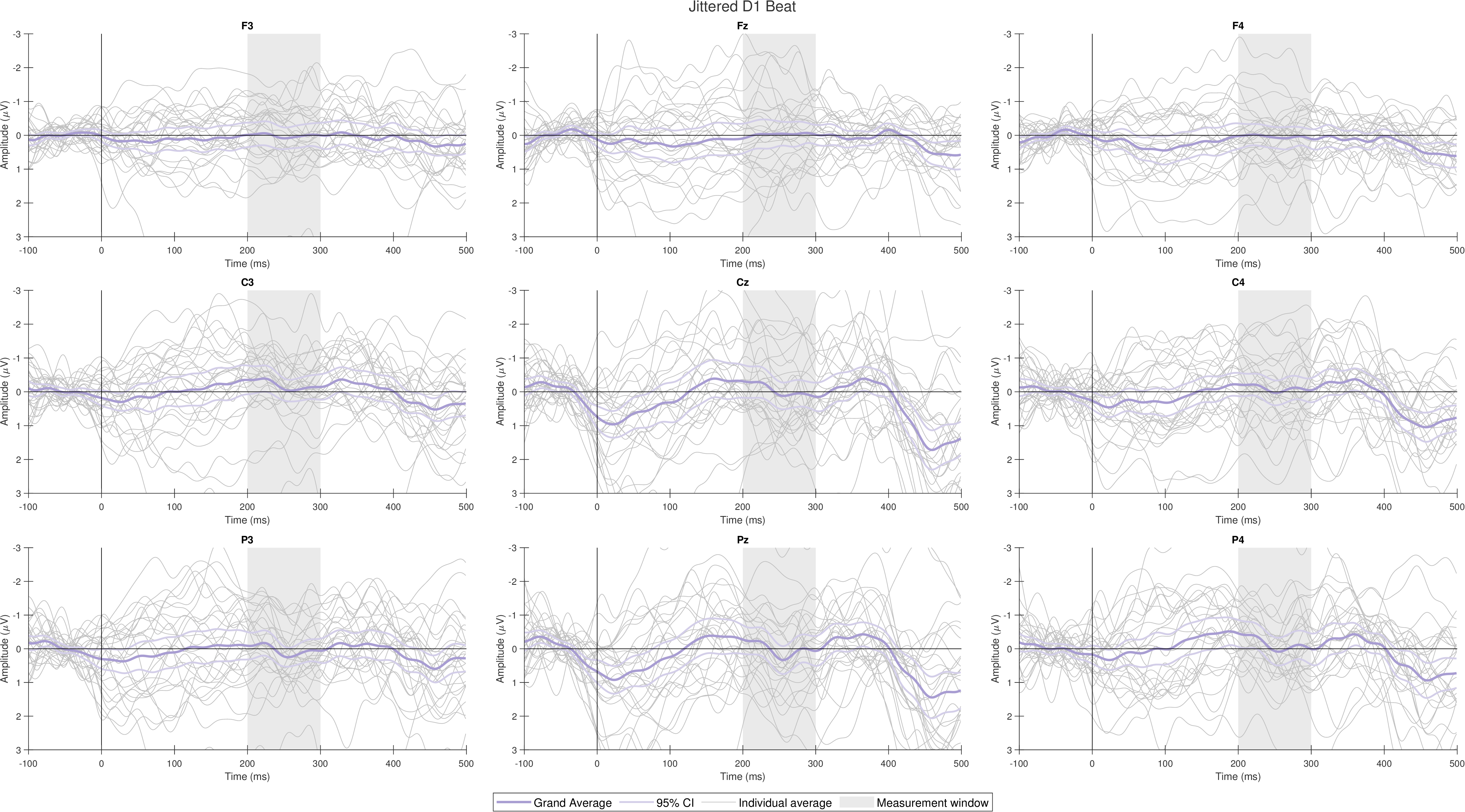

**Supplementary Figure 13**

Individual difference waves (grey) and grand average wave with 95% confidence intervals (color), Jittered D2 Offbeat

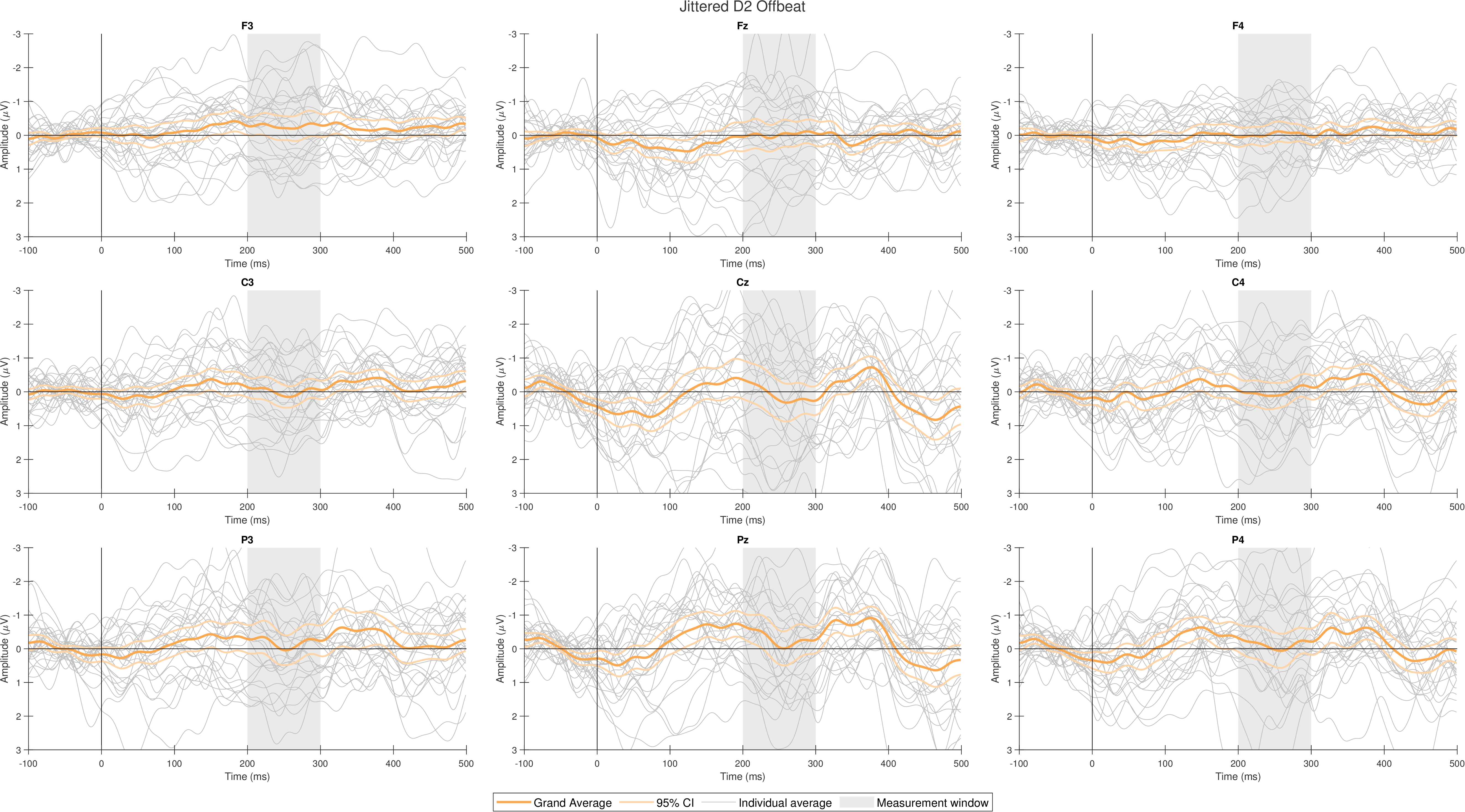

**Supplementary Figure 14**

Individual difference waves (grey) and grand average wave with 95% confidence intervals (color), Jittered S1 Beat

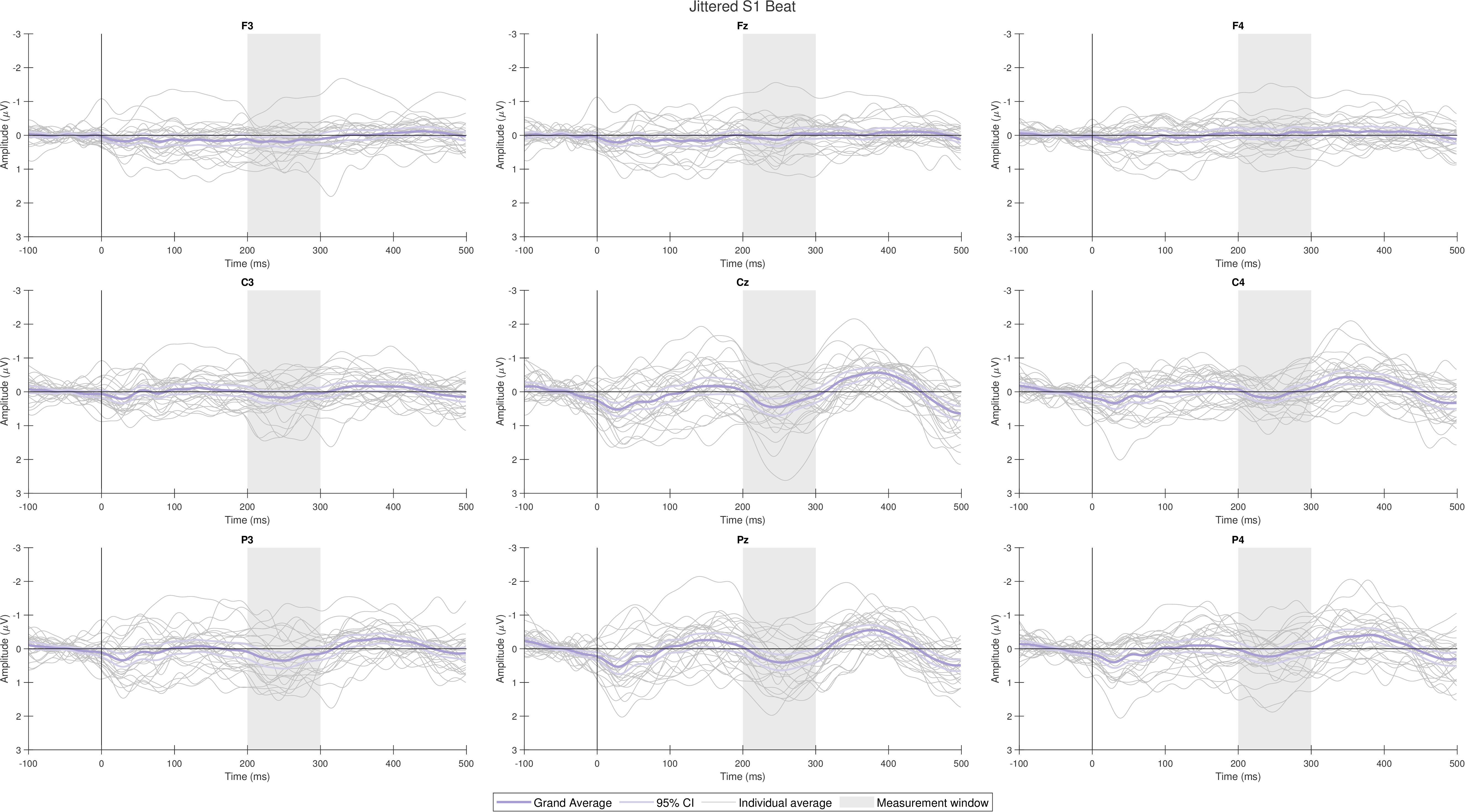

**Supplementary Figure 15**

Individual difference waves (grey) and grand average wave with 95% confidence intervals (color), Jittered S2 Offbeat

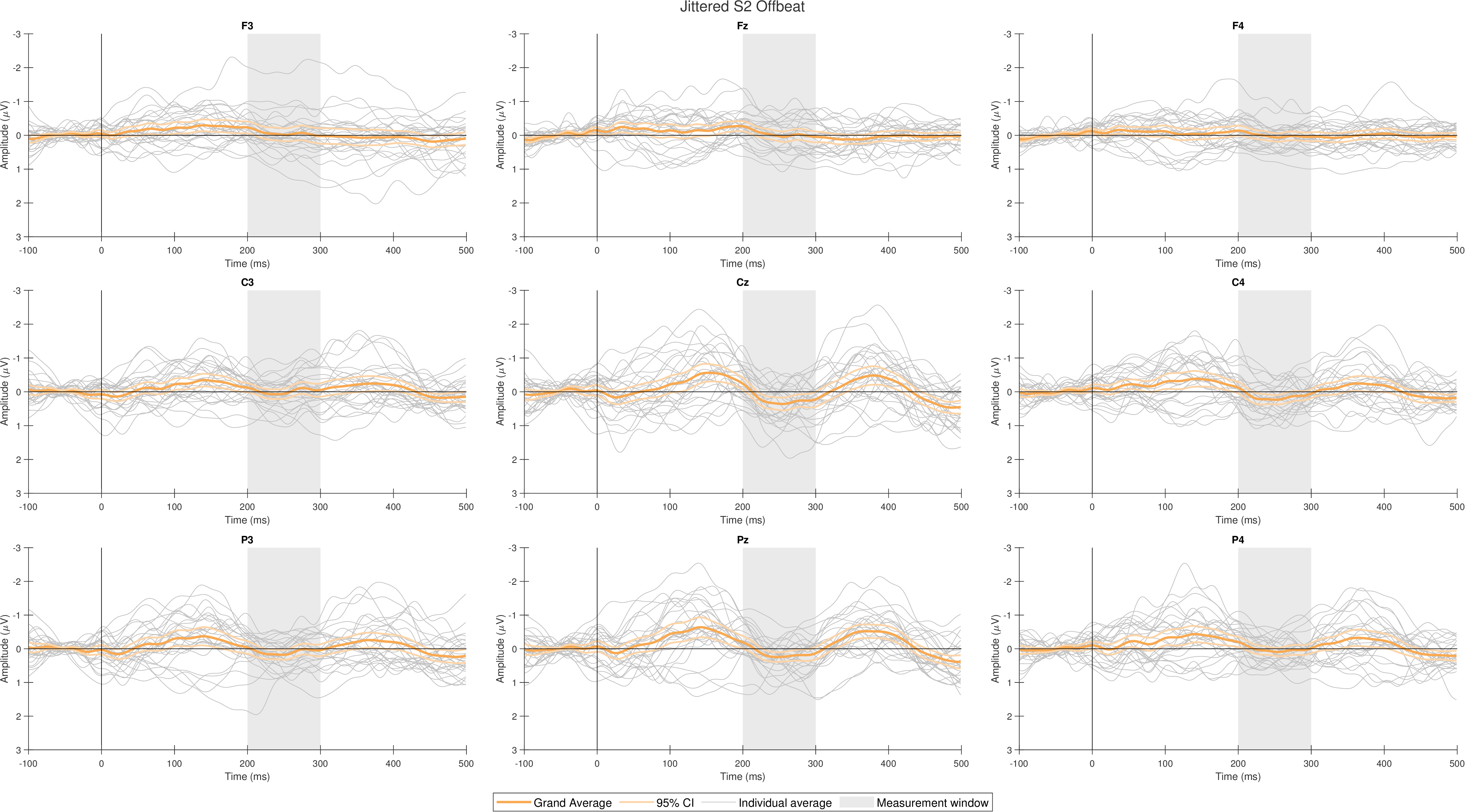

**Output of JASP Software**

The Bayesian t-test for difference between Beat and Offbeat MMR amplitudes

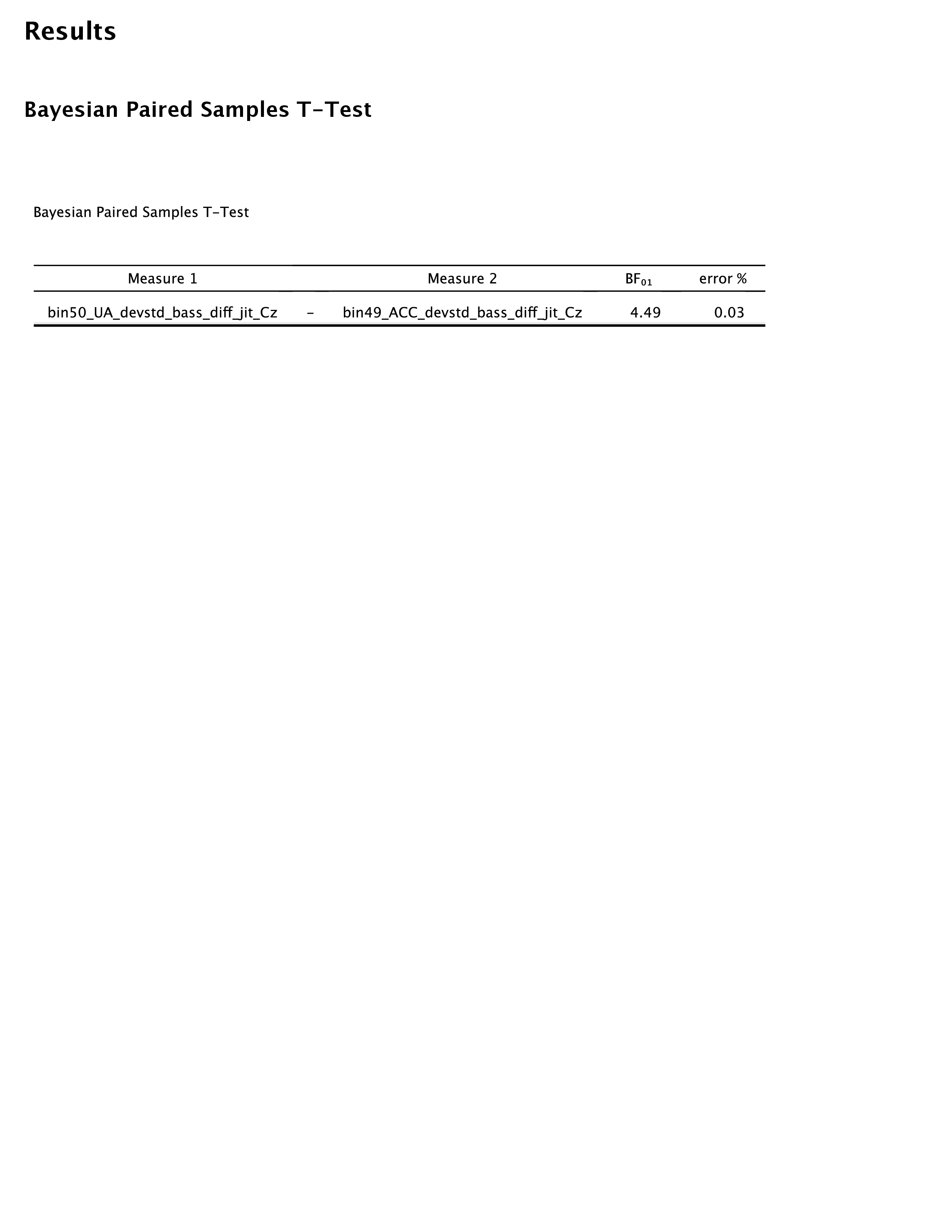

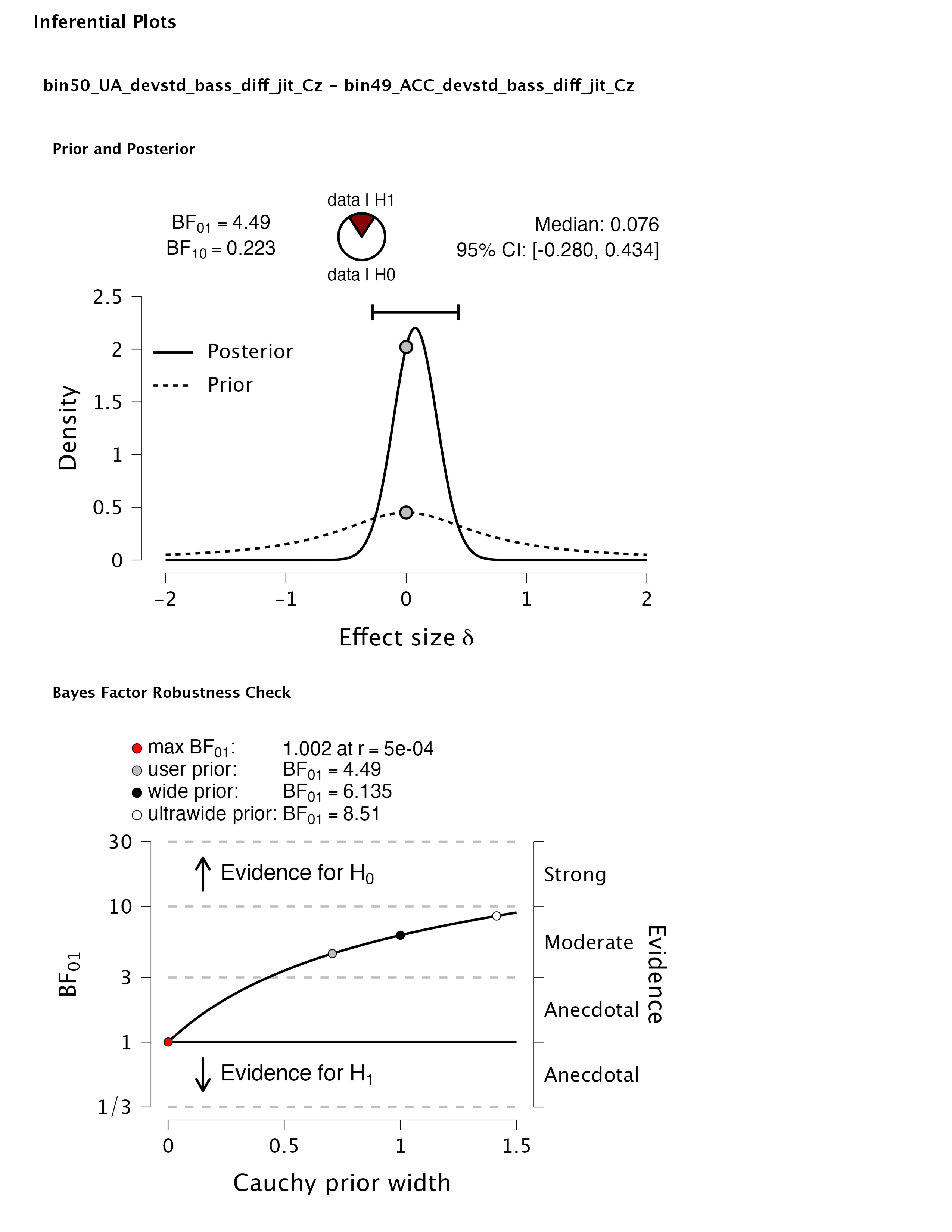

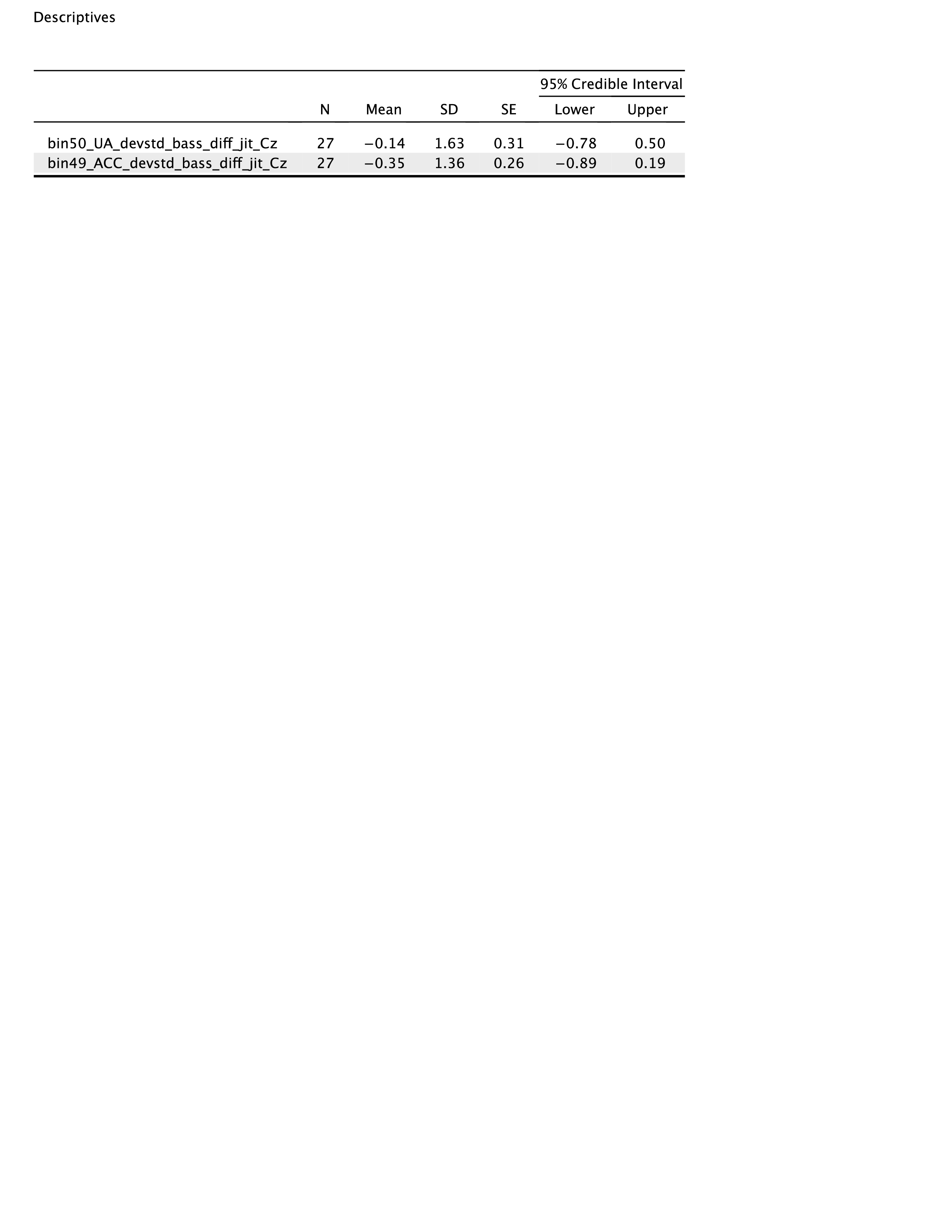
